# Multi-omics profiling, *in vitro* and *in vivo* enhancer assays dissect the *cis*-regulatory mechanisms underlying North Carolina macular dystrophy, a retinal enhanceropathy

**DOI:** 10.1101/2022.03.08.481329

**Authors:** Stijn Van de Sompele, Kent W. Small, Munevver Burcu Cicekdal, Víctor López Soriano, Eva D’haene, Fadi S. Shaya, Steven Agemy, Thijs Van der Snickt, Alfredo Dueñas Rey, Toon Rosseel, Mattias Van Heetvelde, Sarah Vergult, Irina Balikova, Arthur A. Bergen, Camiel J. F. Boon, Julie De Zaeytijd, Chris F. Inglehearn, Bohdan Kousal, Bart P. Leroy, Carlo Rivolta, Veronika Vaclavik, Jenneke van den Ende, Mary J. van Schooneveld, José Luis Gómez-Skarmeta, Juan J. Tena, Juan R. Martinez-Morales, Petra Liskova, Kris Vleminckx, Elfride De Baere

**Author notes:** Correspondence to: Elfride De Baere, MD, PhD, Department of Biomolecular Medicine, Ghent University, Ghent, Belgium; Center for Medical Genetics, Ghent University Hospital, Ghent, Belgium.

## Abstract

North Carolina macular dystrophy (NCMD) is a rare autosomal dominant disease affecting macular development. The disease is caused by non-coding single nucleotide variants (SNVs) in two hotspot regions near *PRDM13* and by duplications in two distinct chromosomal loci, overlapping DNase I hypersensitive sites near either *PRDM13* or *IRX1*.

To unravel the mechanisms by which these variants cause disease, we first established a genome-wide multi-omics retinal database, RegRet. Integration of UMI-4C profiles we generated on adult human retina then allowed fine-mapping of the interactions of the *PRDM13* and *IRX1* gene promoters, and the identification of eighteen candidate *cis*-regulatory elements (cCREs), the activity of which was investigated by luciferase and *Xenopus* enhancer assays.

Next, luciferase assays showed that the non-coding SNVs located in the two hotspot regions of *PRDM13* affect cCRE activity, including two novel NCMD-associated non-coding SNVs that we identified. Interestingly, the cCRE containing one of these SNVs was shown to interact with the *PRDM13* promoter, demonstrated *in vivo* activity in *Xenopus*, and is active at the developmental stage when progenitor cells of the central retina exit mitosis, putting forward this region as a *PRDM13* enhancer.

Finally, mining of single-cell transcriptional data of embryonic and adult retina revealed the highest expression of *PRDM13* and *IRX1* when amacrine cells start to synapse with retinal ganglion cells, supporting the hypothesis that altered *PRDM13* or *IRX1* expression impairs interactions between these cells during retinogenesis.

Overall, this study gained insight into the *cis*-regulatory mechanisms of NCMD and supports that this condition is a retinal enhanceropathy.

**Graphical abstract:** 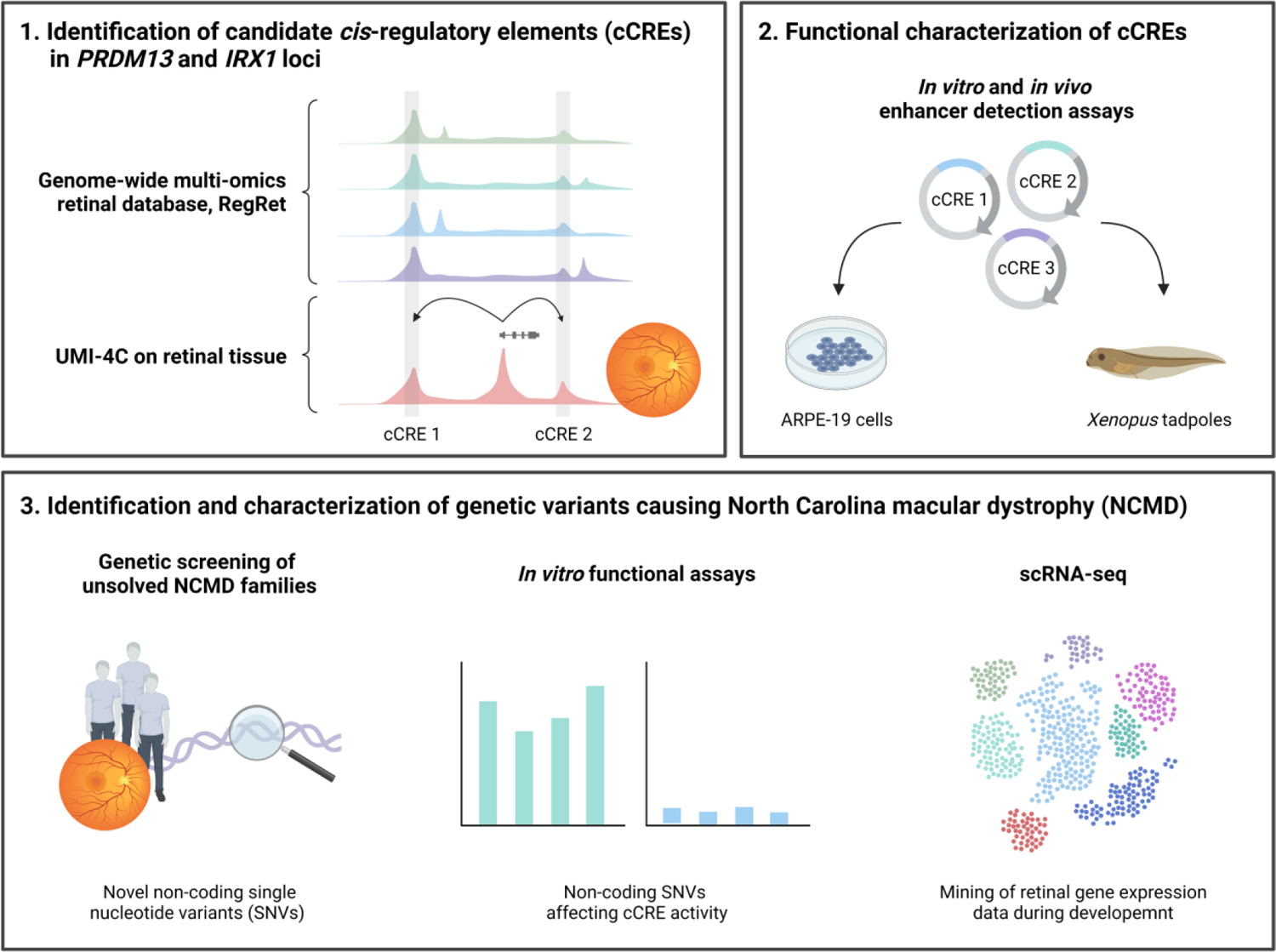

## Introduction

The term “missing heritability”, introduced by Brendan Maher in 2008, refers to the fact that the underlying genetic cause of many complex traits and diseases remains unexplained.^1, 2^ Although many technological advances in the field of genomics took place during the past decade, large-scale exome sequencing still has a limited diagnostic yield ranging from ∼30-60%, depending on the condition being investigated.^3–5^ It was estimated that 85% of mutations with large effects on disease-related traits can be identified in the protein-coding regions. Nevertheless, this number may result from the historical observational bias towards the approximately 1% coding part of the human genome, partly due to the fact that a significant portion of the non-coding genome still lacks proper functional annotation.^6^ In the last couple of years, however, with the continuously reducing cost of next-generation sequencing and the improvements of complementing analysis tools and algorithms, large-scale initiatives have contributed to the better understanding of the human genome and its different functional regions. According to the analysis of the Encyclopedia of DNA Elements (ENCODE) project, based on chromatin accessibility, transcriptional activity, and DNA methylation data, about 80% of the human genome may have a functional significance.^7^ The majority of the identified functional elements, including gene promoters, enhancers, silencers, and insulators, appeared to be involved in spatiotemporal gene expression. Control through these *cis*-regulatory elements (CREs) results from the presence of transcription factor binding sites (TFBS) for tissue- and time-specific transcription factors, which mediate the activation or repression of transcription.^8^ It was also discovered that the human genome contains non-coding elements with an extremely high degree of sequence conservation across multiple vertebrate species. These are called ultra-conserved non-coding elements (UCNEs) and cluster in genomic regions associated with genes known to have a role during development.^9, 10^ Although studies in transgenic animals have shown that some of these elements function as tissue-specific enhancers during developmental processes, the functional significance of others remains to be ascertained.^11, 12^ To facilitate communication between CREs and target gene promoters, the genome is organized into well-defined three-dimensional domains, called topologically associating domains (TADs). This form of genome architecture, which is conserved across cell types, allows long-range interactions between CREs and their target genes within the same TAD, while insulating regulatory activity from neighboring TADs.^13^ Whole genome sequencing (WGS) has revealed structural variants (SVs) that alter the copy number of CREs or disrupt TAD boundaries, resulting in modified three-dimensional chromatin architecture.^14^ Moreover, it has also been shown that single nucleotide variants (SNVs) residing in non-coding CREs could explain a certain fraction of the missing heritability, with the phenotypic consequences usually resulting from alterations in gene expression.^15–18^ Thus far, an increasing number of non-coding variants have been linked to inherited retinal diseases (IRDs), the majority of them affecting *cis*-acting splicing.^19–21^ A well-known example is a deep-intronic mutation in *CEP290*, accounting for ∼20% of congenital blindness due to Leber congenital amaurosis in Northwestern Europe.^22^ In 2016, it was shown that North Carolina macular dystrophy (NCMD, MIM: 136550) is caused by non-coding SNVs near *PRDM13* and duplications overlapping with DNase I hypersensitive sites near either *PRDM13* or *IRX1*, likely dysregulating these transcription factor-encoding genes.^23^

NCMD is a rare autosomal dominant disorder that affects the development of the macula as first noted by Small *et al.* ^23–25^ This completely penetrant maculopathy is present at birth, shows no progression, is characterized by variable expressivity (grade 1 to 3), even among family members, and usually affects both eyes symmetrically.^24, 25^ Grade 1 is characterized by a few small yellow drusen-like lesions within the fovea, which form larger confluent lesions in grade 2. In both grades, patients typically have no or mild impairment of central vision. In grade 3, the fundus has a striking appearance characterized by severe central colobomatous-like chorioretinal atrophy, resulting in mild to moderately impaired vision.^24, 25^ Although NCMD is generally considered as non-progressive, visual deterioration can be observed as a result of complications associated with choroidal neovascularization.^26^ The disease was first described in 1971 upon the clinical investigation of a family from North Carolina,^27^ and twenty years later, NCMD was mapped to chromosome 6q14-q16 (MCDR1 locus, MIM 136550).^28^ During the following decades, linkage analysis was performed in many additional families giving a cumulative logarithm of the odds (LOD) score greater than 40.^29^ Genetic heterogeneity was subsequently found with linkage to 5p15-p13 (MCDR3 locus, MIM 608850) in a single Danish family.^30^ Since no mutations were found in the protein coding regions in both loci, Small *et al.* used WGS which eventually uncovered the first pathogenic non-coding variants underlying NCMD in 2016.^23^

To date, fourteen NCMD-associated genetic variants have been reported, of which eleven are located in the MCDR1 locus and three in the MCDR3 locus. In particular, six pathogenic heterozygous SNVs have been identified in the MCDR1 locus, which cluster together in two distinct hotspots upstream of the *PRDM13* gene.^23, 31, 32^ In addition, five heterozygous tandem duplications have been reported in the same locus, encompassing the *PRDM13* gene and a shared non-coding region (∼44 kb) upstream of the gene, containing several putative CREs.^23, 33–36^ The three other NCMD-associated genetic variants are heterozygous tandem duplications located in the MCDR3 locus. These share a ∼39 kb non-coding region in a gene desert downstream of the *IRX1* gene, while only one out of three duplications encompasses the coding region of *IRX1*.^23, 26^ Interestingly, the shared duplicated region in MCDR3 contains putative CREs active during a definite period of retinal development, as well as a UCNE of which the function is yet unexplored.^9, 26^ Both the *PRDM13* and *IRX1* gene have been demonstrated to have a role during retinal development. More specifically, *PRDM13* encodes a transcription factor characterized by an N-terminal PR (PRDI-BF1 and RIZ1 homology) domain with histone methyltransferase activity, followed by four zinc-finger domains, responsible for protein-protein and protein-DNA binding. ^37^ In the retina, PRDM13 acts as a cell type specification factor downstream of PTF1A, playing an essential role in the development and diversification of specific subsets of amacrine cells. Based on their morphological, neurochemical, and physiological features, more than 30 subtypes of amacrine cells can be defined, each having a distinct set of functions. Most PRDM13-positive amacrine cells use GABA or glycine as a neurotransmitter and adapt visual sensitivity to changing contrast and spatiotemporal frequency. As a consequence, *Prdm13* knockout in the mouse significantly reduces this amacrine cell subtype.^38^ On the other hand, most vertebrate species possess six Iroquois (*IRX*) genes, located in two paralog clusters. These genes encode important developmental homeobox transcription factors, with key roles in regulating gene expression during pattern formation and nervous system development.^39, 40^ In mice, all six *Irx* genes are expressed in the inner layers of the neuroretina during early retinogenesis and *Irx1* is required for the neurogenesis of the inner and outer nuclear layer of the retina. Also in zebrafish, *irx1a* knockdown negatively affects the proper differentiation of amacrine, bipolar, photoreceptor, and Müller glial cells.^41, 42^

In this study, we aimed to provide insight into the *cis*-regulatory mechanisms of NCMD by integrated multi-omics profiling of the human retina, followed by *in vitro* and *in vivo* enhancer assays of candidate *cis*-regulatory elements (cCREs) and NCMD-associated (likely) pathogenic SNVs. Additionally, we expanded the mutational spectrum of NCMD by the identification of novel (likely) pathogenic SNVs in the *PRDM13* region.

## Material and methods

### Generation of an integrated retinal multi-omics database RegRet

In order to gain more insight into the regulatory landscapes of the two NCMD-associated disease loci, multiple publicly available tissue-specific datasets were collected and integrated into a genome-wide database. This database contains multiple chromatin immunoprecipitation sequencing (ChIP-seq) datasets of histone modifications and retinal transcription factors,^43^ assay for transposase-accessible chromatin sequencing (ATAC-seq) data,^43, 44^ and RNA-seq data,^43, 45^ all generated in adult human retina. Moreover, DNase-seq data of embryonic retina at five different stages,^46^ and high-throughput chromosome conformation capture (Hi-C) data from human retinal organoids were included.^47^

### Tissue preparation and nuclei isolation

To fine-map the chromatin interactions within the *PRDM13* and *IRX1* loci in relevant tissue, we first obtained eyes from two healthy human *post-mortem* cornea donors through the Tissue Bank of the Ghent University Hospital under ethical approval of the Ethics Committee of the Ghent University Hospital (2018/1072, B670201837286). Both donors (a 47- and 70-year-old male) did not have a history or clinical evidence of retinal disease. The eyes were transported in CO_2_ Independent Medium (Gibco). Following dissection, retinas were processed according to the protocol of Matelot *et al.* (2016).^48^ Briefly, retinas were treated with a 12.5% collagenase solution and forced through a 40 µm cell strainer, followed by crosslinking using a 2% formaldehyde solution. The crosslinking reaction was quenched by adding a cold 1 M glycine solution on ice. Cell suspensions were then centrifuged and washed with PBS. Cell lysis was performed using a 50 mM Tris-HCl, 150 mM NaCl, 5 mM EDTA, 0.5% NP-40, 1% Triton X-100 lysis buffer containing a complete protease inhibitor cocktail. Next, the lysates were centrifuged to obtain a pellet of nuclei. After a washing step with PBS, the nuclei were aliquoted per 10 million, snap frozen as a pellet in liquid nitrogen, and stored at −80°C. The maximum processing time between enucleation and preservation of nuclei was 24 h.

### RNA isolation, cDNA conversion, qPCR

After retina dissection and before continuing with nuclei isolation, a small fraction of each of the collected retinas was preserved in RNAlater (Qiagen). The tissues were thoroughly homogenized in the presence of TRIzol (Invitrogen). Next, total RNA was obtained through phenol-chloroform extraction. A DNase treatment was performed on-column using the RNeasy mini kit (Qiagen). The iScript cDNA synthesis kit (Bio-Rad) was used for RNA to cDNA conversion and subsequent qPCR analysis was performed using the SYBR Green Master Mix (Bio-Rad) and primers for retina-specific genes and controls. Expression values were analyzed in qbase+ (Biogazelle). Primer sequences can be found in Table S1.

### Generation of UMI-4C sequencing libraries

The dissected retinas were then subjected to unique molecular identifier chromosome conformation capture (UMI-4C) sequencing to identify the chromatin interactions within the two NCMD-associated disease loci. As a first step, libraries were generated according to the protocol of Schwartzman *et al.* (2016).^49^ For every library, the crosslinked DNA from 10 million nuclei was first digested overnight with 400 U *Dpn*II (NEB) at 37°C. Next, samples were diluted one in two before adding 4,000 U T4 DNA ligase (NEB) for overnight incubation at 16°C. Digestion and ligation efficiency were evaluated via agarose gel electrophoresis. DNA was then de-crosslinked with proteinase K (BIOzymTC). After overnight incubation at 65°C, the resulting 3C templates were purified using AMPure XP beads (Agencourt) and 4 µg of each template was sheared on a Covaris M220 focused-ultrasonicator (Covaris) to achieve 300 bp DNA fragments. The obtained fragments were library prepped using NEBNext Ultra II DNA Library Prep Kit (NEB) for Illumina. DNA shearing and library prep efficiency were evaluated on a Fragment Analyzer (Agilent Technologies). Next, a ligation-mediated nested PCR was conducted with an upstream (US) forward primer hybridizing to the selected genomic region of interest (i.e. the viewpoint), a universal reverse primer, 100 ng of library as input, and the KAPA2G Robust HotStart ReadyMix (Roche). After purification using AMPure XP beads (Agencourt), the resulting PCR product was used for the second PCR step, using a downstream (DS) forward primer and the same universal reverse primer. Primer sequences for the different viewpoints can be found in Table S2. After purification, final library composition was evaluated on a Fragment Analyzer (Agilent Technologies).

### UMI-4C sequencing and data-analysis

UMI-4C libraries were multiplexed in equimolar ratios and sequenced on the Illumina NovaSeq 6000 platform at 150 bp paired-end. Sequenced reads were demultiplexed based on their barcodes and their DS primer. UMI-4C data were then analyzed as described in Schwartzman *et al.* (2016) using the umi4cpackage in R (https://bitbucket.org/tanaylab/umi4cpackage).^49^ Briefly, reads containing the primer sequence were first tested for a match with the pad sequence. Reads lacking this match were filtered out. After quality trimming, the restriction fragment ends, representing a potentially informative ligation partner, were mapped to the human genome using Bowtie2.^50^ PCR duplicates were removed based on the UMI sequences, being the last ten bases of the 3′ end of the read pair. Next, restriction fragment ends mapped to genomic coordinates within less than 1,500 bp of the viewpoint were considered as nondigested products and were excluded from the analysis. Contact intensity profiles around the viewpoint were constructed by extracting the number of ligations for all restriction fragment ends within a genomic window and pooling of this data from both samples. Finally, UMI-4C profiles were normalized and domainograms created.

### Generation of *in vitro* reporter constructs

In order to evaluate the activity of cCREs and the effect of NCMD-associated genetic variants, *in vitro* luciferase assays were set up. Therefore, fourteen non-coding regions of interest were PCR amplified from human genomic DNA (Roche) using the Phusion High-Fidelity PCR Kit (NEB). Resulting products were cloned into the pGL4.23 luciferase reporter vector (Promega) by restriction-ligation cloning. The recombinant vectors were amplified in One Shot TOP10 Chemically Competent *E. coli* cells (Invitrogen) and purified using a QIAprep Spin Miniprep Kit (Qiagen). For the generation of the three reporter constructs containing duplicated regions of interest, a second round of restriction-ligation cloning was performed using different restriction enzymes. SNVs were inserted into the regions of interest using the Q5 Site-Directed Mutagenesis Kit (NEB) and mutation-specific primers were designed with the NEBaseChanger tool. The sequence of all inserts was confirmed by Sanger sequencing using the BigDye Terminator v3.1 kit (Life Technologies). All regions of interest, their respective primer sequences, and the mutagenesis primers can be found in Table S3 & S4.

### Luciferase enhancer assays

The recombinant pGL4.23 luciferase reporter vectors were transfected in equimolar amounts into ARPE-19 cells (ATCC, CRL-2302™) using the Lipofectamine 3000 Reagent Protocol, according to the manufacturer’s instructions. The vectors were co-transfected with the pRL-TK *Renilla* luciferase control reporter vector (Promega) for normalization purposes. After 48 h, cells were lysed and luciferase activity was detected using the Dual&Glo Luciferase Assay System (Promega) in a Glomax 96 Microplate Luminometer (Promega). Each transfection was done in triplicate and each experiment was repeated at least three times to ensure reproducibility. For each well, the ratio of firefly luciferase activity was normalized to *Renilla* luciferase activity. Next, these ratios were normalized to the average ratio of the corresponding control vector. For each SNV, the respective wild-type vector was used as a negative control, while the three vectors containing duplicated regions of interest were compared against their respective vectors containing the same region of interest as a single insert. For the set of fourteen non-coding regions of interest, a vector containing an inactive insert of comparable length was used as a negative control. For each variant or region of interest, the effect on luciferase activity was determined using a linear mixed effects model in R, with the luciferase vector as fixed effect and the biological replicate as random effect. P-values were obtained by likelihood ratio tests of the model with the fixed effect against a null model without this effect.

### Cell culture

ARPE-19 cells (ATCC, CRL-2302) were grown in DMEM:F12 (Gibco, No. 30-2006) supplemented with 10% fetal bovine serum, 1% penicillin-streptomycin, 1% non-essential amino acid solution, and 0.1% amphotericin B. Cells were cultured at 37°C and 5% CO_2_ and tested for mycoplasma contamination prior to use.

### Generation of *in vivo* reporter constructs

To functionally assess the potential *in vivo* activities of five cCREs in the *PRDM13* and *IRX1* loci, enhancer detection assays using SED vectors in frog (*Xenopus*) animal models were set up (Table S5). This *Xenopus* transgenesis compatible version of the fluorescent zebrafish enhancer detection (ZED)^51^ vector utilizes a I-*Sce*I based detection system for assessing enhancer activity in *Xenopus* embryos.^52^ A detailed description of the vector can be found in Figure S1. To generate the *in vivo* reporter constructs, the regions of interest were PCR amplified from human genomic DNA (Roche) using *Taq* polymerase (Thermo Scientific) and cloned into the pCR-GW-TOPO vector (Invitrogen). Using LR Clonase II (Invitrogen), the DNA fragments were then subcloned into the destination SED vector, as previously described.^53^

### Functional characterization of cCREs using enhancer assays in *Xenopus* embryos

All experiments on *Xenopus tropicalis* and albino *Xenopus laevis* were executed in accordance with the guidelines and regulations of Ghent University, Faculty of Sciences, Belgium. Approval (EC2014-089 and EC2017-104) was obtained by the ethical committee of Ghent University, Faculty of Sciences. For both *Xenopus tropicalis* and albino *Xenopus laevis*, females and males were primed with 20 U and 10 U human chorionic gonadotropin (hCG, Pregnyl, Merck), respectively. Natural mating was set-up the next day for *Xenopus tropicalis* and 5 days after priming for albino *Xenopus laevis*, after boosting the females and males with 150 and 100 U hCG, respectively. All the injections took place at the day of mating. Therefore, the I-*Sce*I meganuclease-mediated transgenesis protocol was followed, as previously described for *Xenopus*.^52^ Briefly, a total of 100 ng of SED vector construct was digested with 5 U of I-*Sce*I meganuclease (NEB) at 37°C for 10 minutes in a 10 μL reaction mixture. Fertilized *Xenopus* eggs were obtained and dejellied using established protocols.^54^ Embryos were then injected either unilaterally in the two-cell stage, or in two of the dorsal blastomeres at the four-cell stage, using 1 nL of the reaction mixture. Next, embryos were raised overnight at room temperature and replaced at 25.5°C for the rest of their development. Embryos were screened for enhanced green fluorescent protein (EGFP) reporter expression at different developmental stages using fluorescent microscopy.

### Targeted sequencing and qPCR

To expand the mutational spectrum of NCMD, targeted genetic testing was performed in a cohort of twenty-three unsolved and unrelated index cases with a clinical diagnosis of NCMD. This study was approved by the ethics committee for Ghent University Hospital (2018/1566, B670201938572). The known NCMD-associated SNVs were analyzed on genomic DNA by PCR amplification of the two mutational hotspots, respectively containing the variants referenced as V1-V3, V12 and the V10-V11 variants, followed by Sanger sequencing using the BigDye Terminator v3.1 kit (Life Technologies). For detection of the previously reported tandem duplications, primers specific for the duplication products were designed. Fragment analysis was performed on a Fragment Analyzer (Agilent Technologies) using the PROSize 2.0 software. All primer sequences can be found in Table S6. In the next step, copy number variant analysis was performed for a selection of regions implicated in the known tandem duplications. In particular, the copy number of the coding regions of *PRDM13* and *IRX1*, as well as the shared duplicated region downstream of *IRX1* were evaluated by qPCR using the SYBR Green Master Mix (Bio-Rad). Copy number values were analyzed in qbase+ (Biogazelle). All primer sequences can be found in Table S6.

### Whole genome sequencing (WGS)

Following targeted sequencing and copy number variant analysis, a subset of nine molecularly undiagnosed NCMD index cases was subjected to WGS. Briefly, samples were prepared according to the Illumina TruSeq DNA PCR-free library preparation guide. The library of family F1 was sequenced on the Illumina HiSeq X-Ten platform at 350 bp paired-end, while all other libraries were sequenced on the Illumina NovaSeq 6000 platform at 150 bp paired-end. Reads were mapped to the human genome (hg38) using Isaac aligner (iSAAC-04.18.11.09)^55^ and Strelka (2.9.10)^56^ was used to identify SNVs and short indels. Variants were annotated by SnpEff (v4.3t)^57^ in combination with additional databases, including ESP6500,^58^ ClinVar,^59^ and dbNSFP3.5.^60^ To identify SVs and large indels, the Manta (1.5.0)^61^ software was utilized. At first, the two NCMD-associated disease loci were analyzed for candidate variants. Therefore, SNVs were filtered based on zygosity (heterozygous), Genome Aggregation Database (gnomAD v3.1.2)^62^ population frequency (<0.005%), and genomic location (2 Mb and 4 Mb region around the *PRDM13* and *IRX1* gene, respectively corresponding to the TAD the gene is located in). SVs located in these genomic regions were evaluated using the Database of Genomic Variants (DGV, http://dgv.tcag.ca/dgv/app/home). Next, the coding regions of 290 known retinal disease genes (RetNet panel V5, https://sph.uth.edu/retnet/disease.htm) were assessed for (likely) pathogenic variants, both SNVs and SVs, that could explain a macular phenotype. Data analysis was performed using our in-house developed analysis platform Seqplorer, which incorporates gnomAD population frequencies, *in silico* missense predictions (REVEL, PolyPhen-2, SIFT, CADD, etc.), and splice predictions (MaxEntScan, SpliceRegion). Variants were confirmed by Sanger sequencing using the BigDye Terminator v3.1 kit (Life Technologies) and underwent segregation analysis where possible. All primer sequences can be found in Table S6.

### Clinical evaluation

Ophthalmic examination of the cohort of twenty-three unsolved NCMD families included fundus photography, optical coherence tomography (OCT), and measurement of best-corrected visual acuity (BCVA). In family F1, color vision was tested by means of a Lanthony D-15 saturated and desaturated assay.

### *In silico* assessment of the non-coding SNVs in the mutational hotspots

For the previously reported and novel pathogenic NCMD-associated SNVs in the two mutational hotspots upstream of *PRDM13*, potential effects of the nucleotide changes on TFBS motifs were analyzed using the TRANSFAC software.^63^ Furthermore, an *in silico* assessment was performed using several functional prediction tools for human non-coding regulatory variants, including an integrated non-coding regulatory prediction score, regBase,^64^ as well as several other tools (GenoCanyon,^65^ ncER,^66^ DVAR,^67^ FIRE,^68^ PAFA,^69^ CDTS^70^).

### *In silico* assessment of the NCMD-associated duplication breakpoints

A bioinformatics analysis was performed for the breakpoints of previously reported pathogenic NCMD-associated tandem duplications in the *PRDM13* and *IRX1* loci, as previously described.^71, 72^ In particular, the degree of microhomology (multiple sequence alignment, ClustalW), the presence of repetitive elements (RepeatMasker track UCSC Genome Browser), and the presence of sequence motifs, based on 40 previously described motifs (Fuzznuc) were analyzed on the breakpoint regions of the tandem duplications.^73, 74^

### Data mining in single-cell retinal dataset

Publicly available single-nucleus RNA-seq data of embryonic (53, 59, 74, 78, 113, and 132 days of development) and adult (25, 50, and 54 years old) human retinal cells, generated by Thomas *et al.* (2022), were processed for evaluating *PRDM13* and *IRX1* expression at single-cell level.^75^ Expression matrices derived from nine *post-mortem* donor neural retinal samples (GSE183684) were imported into R (v4.0.5) and processed using the Seurat single-cell analysis package (v4.0).^76^ Pre-processing and quality control was conducted to remove outlier cells. Briefly, we only considered genes with counts in at least three cells and filtered out cells that had unique feature (gene) counts <200 or >9,000 and that expressed >5% mitochondrial counts. The data were then total-count normalized, logarithmized, filtered for highly variable features, and scaled to unit variance. We used the Harmony package to merge all the single-cell data.^77^ After quality control, pre-processing, and merging of all the data, a total of 60,014 retinal cells were kept for subsequent dimensionality reduction, embedding, and clustering. Markers associated with major neural retina cell populations were used to assess *IRX1* and *PRDM13* expression at single-cell level.

## Results

### Establishment of RegRet, a genome-wide multi-omics retinal database

Multiple publicly available retinal datasets from different sources were integrated in a genome-wide multi-omics database. In particular, the database contains ChIP-seq datasets of histone modification H3K27ac from adult human retina, macula, and retinal pigment epithelium (RPE), and H3K4me2 from adult human retinal tissue, both indicative of enhancer activity.^43^ Additional ChIP-seq datasets of retinal (CRX, NRL, OTX2, MEF2D, RORB) and other (CTCF, CREB) transcription factors, generated in adult human retina were also included.^43^ Next, we integrated available ATAC-seq data from both adult whole retina and macula,^43^ as well as ATAC-seq data from adult retina and RPE, where a distinction was made between macular and peripheral regions.^44^ In the context of retinal transcriptomics, quantitative RNA-seq data and newly identified transcripts observed in adult human retina were added,^45^ next to single-nucleus RNA-seq data generated in adult human retina, macula, and RPE.^43^ With regard to retina-specific chromatin interactions, Hi-C data from human retinal organoids was added.^47^ To provide insight into retinal development, DNase-seq data from embryonic retinal tissue at five stages (74, 85, 89, 103, and 125 days) were included.^46^ Apart from the retina-specific datasets, multiple non-retinal tracks were added: UCNEs and ultraconserved gene regulatory blocks (UGRBs),^9^ H3K27ac, H3K4me1, and H3K4me3 marks in seven cell lines (ENCODE Regulation), DNase I hypersensitive site peak clusters in 95 cell lines (ENCODE Regulation), ORegAnno elements,^78^ ENCODE cCREs, RefSeq functional elements, GeneHancer regulatory elements and gene interactions, GeneCards gene transcription start sites, and Micro-C chromatin structure in H1-hESC. This genome-wide database, called RegRet (http://genome.ucsc.edu/s/stvdsomp/RegRet), is accessible via the UCSC Genome Browser and served as a basis for the subsequent locus-specific research of the NCMD-associated disease regions. Furthermore, it is also of use for studies of other retinal disease-associated loci.

### UMI-4C profiling on human adult retina reveals cCREs in the *PRDM13* and *IRX1* loci

To identify the non-coding regions in the two known disease loci that interact with the *PRDM13* and *IRX1* promoters, we performed chromosome conformation capture, in particular UMI-4C, on retinas of adult human donor eyes. After dissection of the eyes, the purity of the retinal tissue was first demonstrated by expression analysis of several retina- and RPE-specific genes (Figure S2). Next, UMI-4C was performed on the cross-linked retinal nuclei, using the *PRDM13* and *IRX1* promoter regions as bait sequences. The numbers of raw, mapped, and unique reads for each viewpoint can be found in Table S7. For both genes, UMI-4C profiles indicated that promoter interactions were spread across but limited to their respective TAD (Figure 1). These profiles were then integrated into the retina-specific multi-omics RegRet database, which enabled the identification of cCREs in the two NCMD-associated disease loci. In particular, cCREs were defined as non-coding regions interacting with the *PRDM13* or *IRX1* gene promoter while demonstrating overlap with peaks of epigenomics datasets or containing a UCNE. Although, based on our UMI-4C experiments, the first mutational hotspot upstream of *PRDM13* and the shared duplicated region downstream of *IRX1* do not exhibit interactions with the *PRDM13* or *IRX1* gene promoter in the adult retina, these regions were nevertheless considered as cCREs in the course of this study (PRDM13_cCRE3, IRX1_cCRE10). Reason for this is that both regions are important for NCMD disease pathogenesis, while overlapping with DNase I hypersensitive sites during retinal development and with ChIP-seq peaks of retinal transcription factors or a UCNE, respectively. This brings the total number of identified cCREs to eight and ten for the *PRDM13* and *IRX1* locus, respectively (Table 1 & Figure S3).

**Figure 1.**
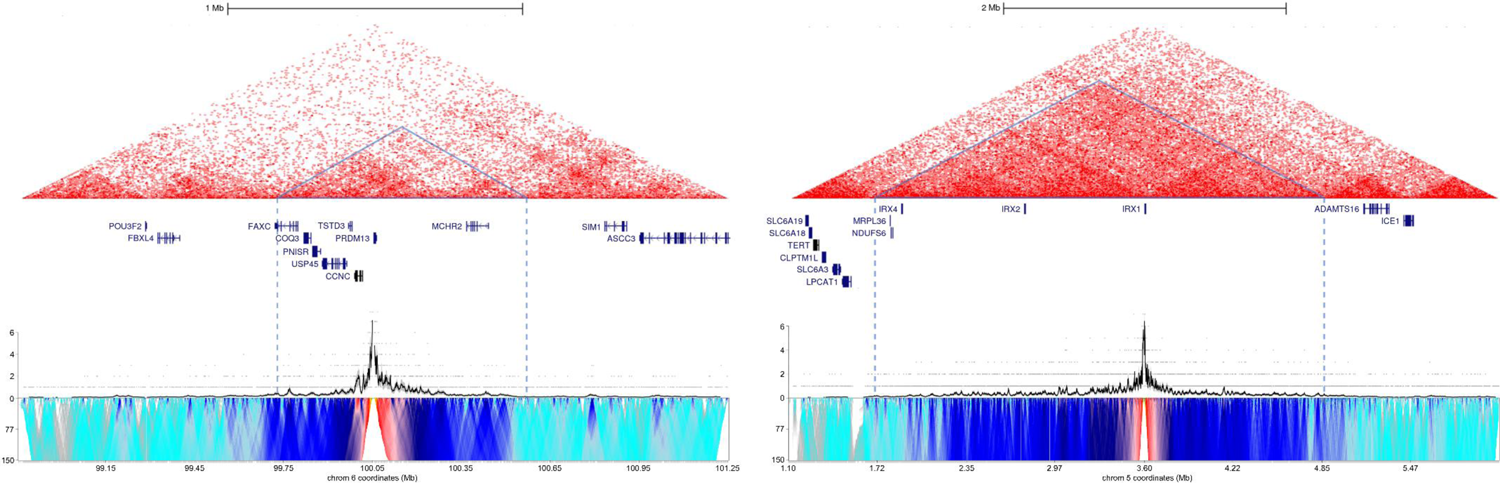
Integration of the generated UMI-4C data with publicly available Hi-C data. The UMI-4C interaction frequency profiles and domainograms (bottom) for the *PRDM13* promoter (left) and *IRX1* promoter (right) viewpoints were integrated with Hi-C data from control human retinal organoids (top), demonstrating promoter interactions within the respective TADs.^47^ Topologically associated domains (TADs) are indicated by blue triangles. Chromosome coordinates are in hg38 annotation.

**Table 1.** Overview of cCREs in the *PRDM13* and *IRX1* loci. The cCREs are obtained after integration of the UMI-4C profiles in the retina-specific multi-omics database (Figure S3). For each cCRE, it is indicated which characteristics contributed to their identification. The two right columns indicate whether cCRE activity was assessed using *in vitro* luciferase assays or *in vivo* enhancer detection assays, as well as the outcome of these experiments. cCRE: candidate CRE, DHS: DNase I hypersensitive site, ChIP-seq: chromatin immunoprecipitation sequencing,, UCNE: ultraconserved non-coding element.

| cCRE | Coordinates (hg38) | cCRE characteristics | *In vitro* luciferase assay | *In vivo* enhancer detection assay |
| --- | --- | --- | --- | --- |
| *PRDM13* locus |  |  |  |  |
| PRDM13_cCRE1 | chr6:99,588,208-99,589,239 | UMI-4C peak, DHS, ChIP-seq (H3K4me2, CRX, NRL, OTX2) | Not significant | Eye and brain |
| PRDM13_cCRE2 | chr6:99,590,566-99,591,414 | UMI-4C peak, DHS | / | / |
| PRDM13_cCRE3 | chr6:99,592,841-99,593,364 | First mutational hotspot, DHS, ChIP peak (CRX, OTX2) | Positive | No activity |
| PRDM13_cCRE4 | chr6:99,595,852-99,596,746 | UMI-4C peak, DHS | / | / |
| PRDM13_cCRE5 | chr6:99,598,651-99,599,224 | Second mutational hotspot, UMI-4C peak, DHS | Not significant | Eye and brain |
| PRDM13_cCRE6 | chr6:99,617,828-99,619,867 | UMI-4C peak, DHS, ChIP-seq (H3K27me2) | Positive | / |
| PRDM13_cCRE7 | chr6:99,645,605-99,647,017 | UMI-4C peak, DHS, highly conserved region | / | / |
| PRDM13_cCRE8 | chr6:99,749,318-99,752,123 | UMI-4C peak, DHS, ChIP-seq (H3K4me2, H3K27ac), UCNE | Not significant | / |
| *IRX1* locus |  |  |  |  |
| IRX1_cCRE1 | chr5:3,224,667-3,227,420 | UMI-4C peak, DHS, ChIP-seq (H3K4me2, H3K27ac), UCNE | Positive | / |
| IRX1_cCRE2 | chr5:3,426,706-3,428,595 | UMI-4C peak, DHS, UCNE | Positive | / |
| IRX1_cCRE3 | chr5:3,488,008-3,490,747 | UMI-4C peak, DHS, ChIP-seq (H3K27ac), UCNE | Positive | / |
| IRX1_cCRE4 | chr5:3,529,207-3,531,844 | UMI-4C peak, UCNE | Negative | / |
| IRX1_cCRE5 | chr5:3,620,470-3,622,600 | UMI-4C peak, DHS | Negative | / |
| IRX1_cCRE6 | chr5:3,628,385-3,630,819 | UMI-4C peak, DHS, conserved region | Not significant | / |
| IRX1_cCRE7 | chr5:3,649,694-3,651,723 | UMI-4C peak, DHS, conserved region | Positive | Eye, brain, neural crest |
| IRX1_cCRE8 | chr5:3,729,667-3,732,532 | UMI-4C peak, DHS, conserved region | Positive | / |
| IRX1_cCRE9 | chr5:3,786,388-3,788,778 | UMI-4C peak, DHS, conserved region | / | / |
| IRX1_cCRE10 | chr5:4,424,583-4,425,559 | Shared duplicated region, DHS, UCNE | Not significant | Eye and brain |

One particularly interesting cCRE was identified upstream of the *PRDM13* promoter (PRDM13_cCRE1). There, the UMI-4C data demonstrated a distinct interaction with the *PRDM13* promoter, in addition to an overlap with DNase-seq and ATAC-seq peaks, as well as with ChIP-seq profiles of specific histone marks indicative of enhancer activity (H3K27ac, H3K4me2) and ChIP-seq profiles of retinal transcription factors (CRX, NRL, OTX2) (Figure 2). To confirm this interaction, a reverse UMI-4C experiment was conducted using this cCRE as a bait sequence. The resulting UMI-4C profile confirmed the interaction between the CRE and the *PRDM13* promoter, altogether making this region a very likely enhancer of *PRDM13* (Figure 2).

**Figure 2.**
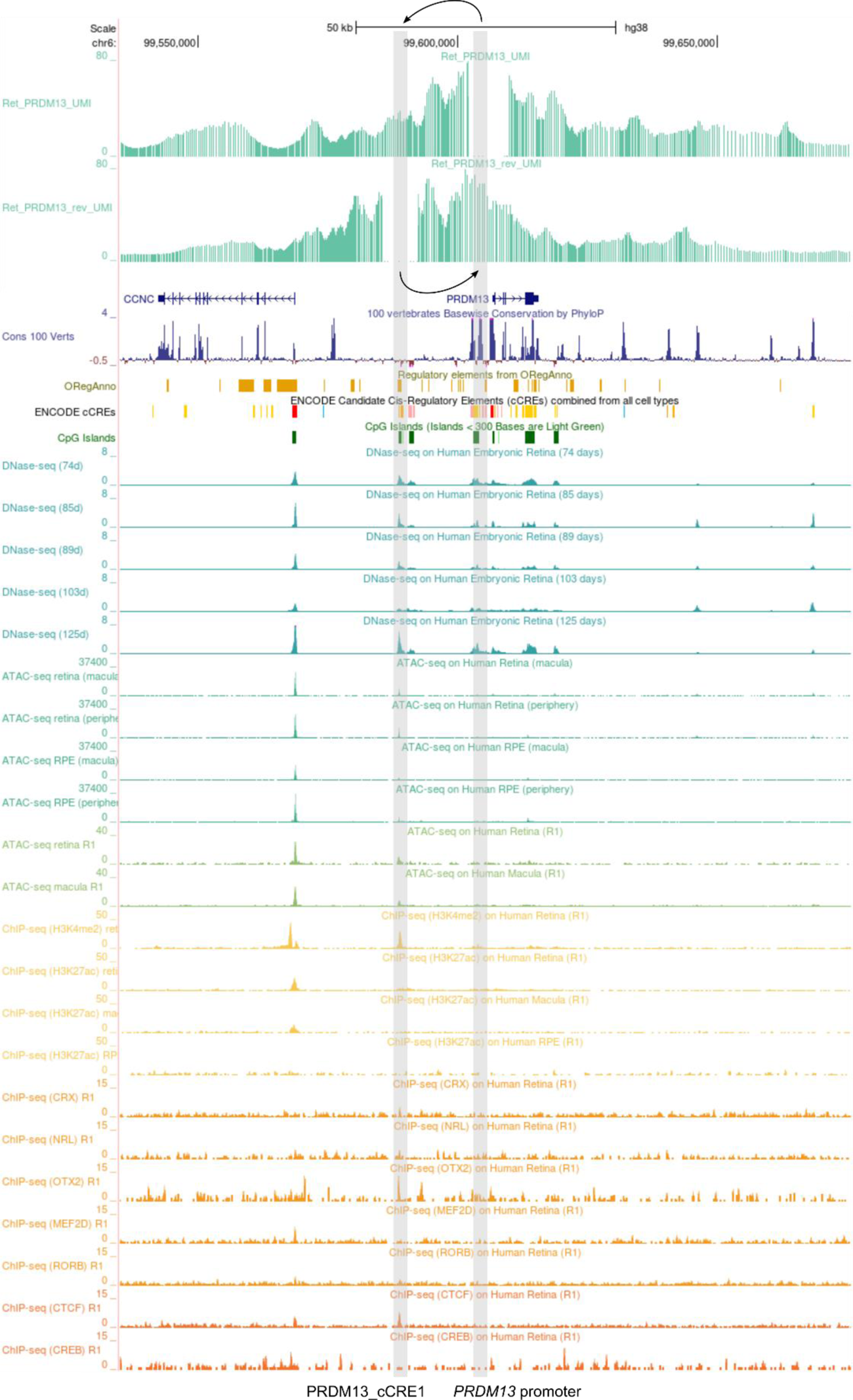
Output from the UMI-4C experiment in the *PRDM13* locus. The UMI-4C data (green) from the *PRDM13* promoter viewpoint (right gray bar) illustrate an interaction with PRDM13_cCRE1 (left gray bar), a non-coding region upstream of the promoter (left arrow). Since underlying epigenomic tracks show an overlap of this region with DNase-seq (turquoise) and ATAC-seq (green) peaks, as well as with ChIP-seq profiles of specific histone marks indicative of enhancer activity (H3K4me2, H3K27ac) (yellow) and ChIP-seq profiles of retinal transcription factors (CRX, NRL, OTX2) (orange), this region is a strong cCRE. The reverse UMI-4C experiment using this cCRE as a viewpoint, results in a peak around the *PRDM13* promoter region (right arrow), confirming this interaction. cCRE: candidate CRE.

A second reverse UMI-4C experiment was performed for the shared duplicated region located in a gene desert ∼800 kb downstream of the *IRX1* gene (IRX1_cCRE10). Although we were not able to identify an interaction using the *IRX1* promoter as a viewpoint, the reverse UMI-4C experiment with IRX1_cCRE10 as a viewpoint suggested an interaction between the two regions (Figure S4).

### *In vitro* enhancer assays show a regulatory effect for cCREs in the *PRDM13* and *IRX1* loci

From the eighteen identified cCREs, a set of fourteen cCREs was selected to determine their *in vitro* enhancer activity. These regions were cloned into separate luciferase reporter vectors (pGL4.23) (Table S3) and a negative control luciferase vector with an inactive insert of comparable length was used as reference. The subsequent luciferase assays in ARPE-19 cells demonstrated that seven of the tested regions were able to significantly increase luciferase reporter levels over the negative control vector (P<0.001), while two regions in the *IRX1* locus resulted in a decrease of luciferase expression (P<0.001) (Figure 3 & Table S8). Remarkably, five regions, some of which had strong *in silico* predictions, did not show a significant difference in luciferase reporter levels compared to the negative control vector (P>0.05).

**Figure 3.**
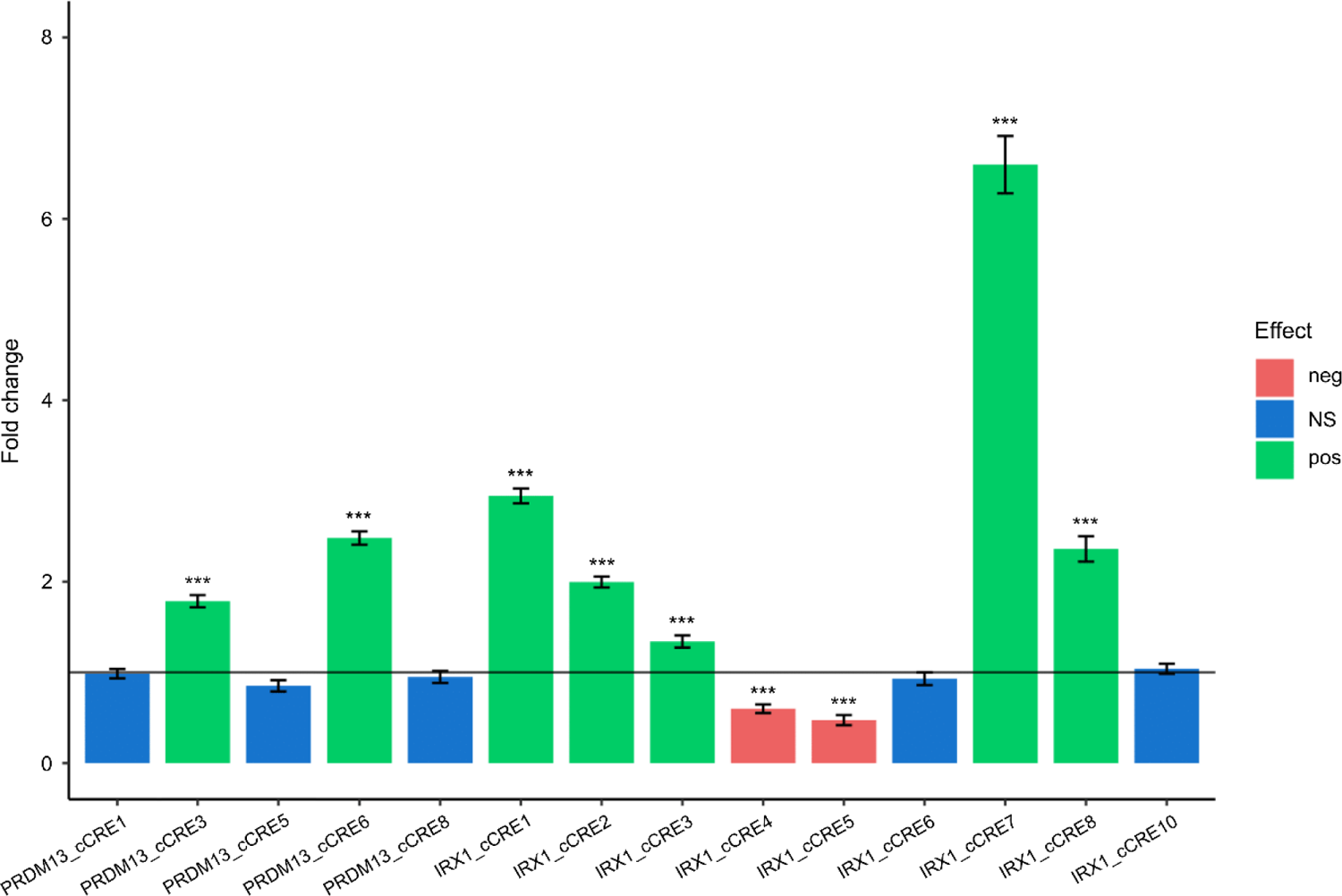
Results from the luciferase assays for the set of fourteen cCREs. The bar plot shows, for each cCRE, the fold change of the luciferase reporter level relative to the level of the negative control luciferase vector (fold change = 1). neg: negative, NS: not significant, pos: positive, ***: P<0.001.

### *In vivo* enhancer assays in *Xenopus* display eye- and brain-specific activity for cCREs in the *PRDM13* and *IRX1* loci

In order to address the activity of cCREs *in vivo*, the five most prominent cCREs were cloned into the SED vector, upstream of a minimal *gata2* promoter element and an EGFP reporter gene. In particular, the cCRE with the highest *in vitro* activity from the *IRX1* locus (IRX1_cCRE7) and the shared duplicated region (IRX1_cCRE10), as well as the cCRE with the strongest epigenomic signatures from the *PRDM13* locus (PRDM13_cCRE1) and the two mutational hotspots (PRDM13_cCRE3, PRDM13_cCRE5) were selected. All constructs were injected unilaterally in two-cell stage *Xenopus tropicalis* embryos, the non-injected side of which acted as an internal negative control. In line with the *irx1* expression in the neural plate in neurula stages of *Xenopus* embryos, the IRX1_cCRE7 construct drove EGFP reporter expression in neural plate and tube in the injected side of the embryos at Nieuwkoop and Faber (NF) stage 15 and 20 (Figure 4A & 4B).^79^ At NF stage 42/43 and 45, EGFP reporter expression was detected in the craniofacial cartilage, a derivative of the cranial neural crest (NC), and in the eye (Figure 4C & 4D). In these transgenic tadpoles, DsRed expression, which in the SED vector is driven by the *Xenopus* cardiac actin promoter, was visible in muscles, thereby demonstrating successful transgene integration (Figure 4E & 4F). When the tadpoles were grown and screened for EGFP reporter expression before metamorphosis at NF stage 55, 13 out of 30 transgenic embryos were found to express EGFP in the eye of the injected side, which was visible through the lens of the dark-pigmented eye (Figure 4G). Meanwhile, no fluorescent signal was observed in the eye on the non-injected side of these tadpoles (Figure 4H). The findings from the other *Xenopus tropicalis* transgenesis experiments are summarized in Table S9.

**Figure 4.**
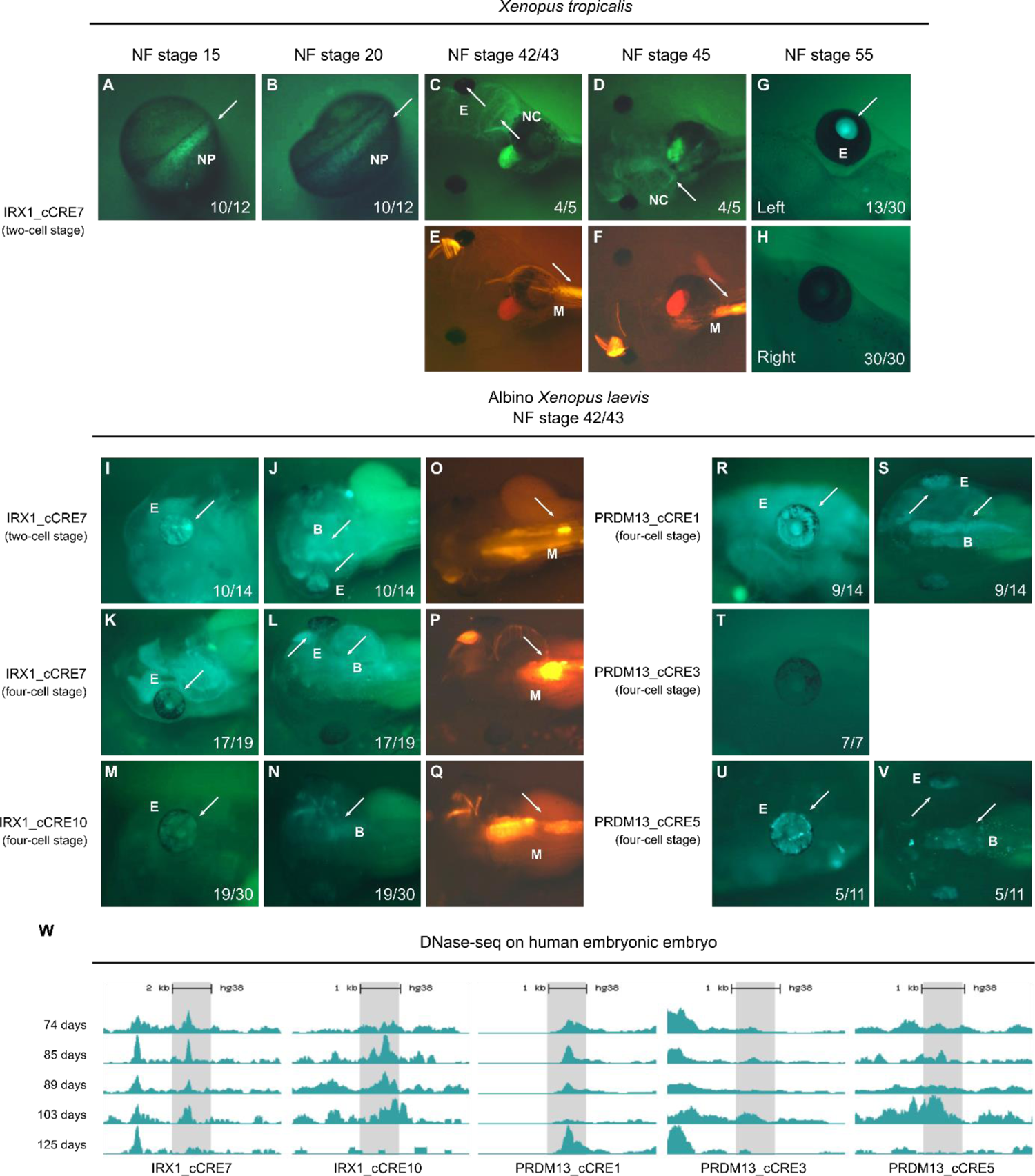
Overview of the results from the *in vivo* transgenic enhancer assays in *Xenopus*. Representative images of EGFP reporter expression in living, transgenic *Xenopus* tadpoles, driven by SED vector reporter constructs containing non-coding regions of interest. The respective *Xenopus* species injected and the Nieuwkoop and Faber (NF) stage shown in the pictures are indicated on top of the images, while the injected construct and the stage of injection are indicated left of the images. For each stage, the number of tadpoles displaying the depicted EGFP reporter expression pattern over the total number of analyzed transgenic tadpoles is given. (**A,B**) Dorsal view of transgenic *Xenopus tropicalis* embryos upon unilateral injection of the reporter construct containing IRX1_cCRE7. EGFP reporter expression was visible at the neural plate and tube (NP) on the injected side, indicated by the arrow, while no expression was observed on the non-injected side. (**C,D**) Ventral view of transgenic tadpoles, displaying EGFP reporter expression in the craniofacial cartilage, a derivative of the neural crest (NC), and the eye (E) on the injected side. (**E,F**) At the same stages, DsRed (positive control) expression was apparent in the myotomes (M) of the tadpoles. (**G,H**) Detailed view of the eye at NF stage 55 indicates that EGFP reporter expression was maintained on the injected (left) side, while no expression was observed in the non-injected (right) side. (**I,J**) Lateral and dorsal view of transgenic albino *Xenopus laevis* tadpoles upon unilateral injection of the reporter construct containing IRX1_cCRE7 in the two-cell stage demonstrated EGFP reporter expression in the eye and brain (B). (**K,L**) Lateral and dorsal view of transgenic tadpoles injected with the same construct, but in two of the dorsal blastomeres at the four-cell stage, also displayed EGFP reporter expression in the eye and brain. (**M,N**) Lateral and dorsal view of transgenic tadpoles introduced with the IRX1_cCRE10 reporter construct demonstrated low EGFP reporter expression in the eye and brain. (**O-Q**) DsRed (positive control) was expressed in the myotomes of the corresponding tadpoles. (**R,S**) Lateral and dorsal view of transgenic albino *Xenopus laevis* tadpoles upon injection of the reporter construct containing the PRDM13_cCRE1 region demonstrated EGFP reporter expression in the eye and brain. (**T**) Lateral view of transgenic tadpoles injected with the first mutational hotspot (PRDM13_cCRE3) showed no EGFP reporter expression in the eye or brain. (**U,V**) Lateral and dorsal view of transgenic tadpoles injected with the second mutational hotspot (PRDM13_cCRE5) demonstrated EGFP reporter expression in the eye and brain. (**W**) For the five cCREs analyzed using *in vivo* enhancer detection assays, DNase-seq profiles generated in human embryonic retinal tissue at five different developmental stages are given. In case of IRX1_cCRE7, IRX1_cCRE10, PRDM13_cCRE1, and PRDM13_cCRE7, open chromatin is observed at/until day 103 of development, while PRDM13_cCRE1 is closed exclusively at this period.

As EGFP reporter expression driven by the SED vector reporter constructs becomes obscured by the pigmentation of the eyes in the tadpole stages, albino *Xenopus laevis* animals were used to obtain embryos with non- or poorly-pigmented eyes. Transgenic albino embryos that were unilaterally injected with the IRX1_cCRE7 reporter construct in the two-cell stage demonstrated EGFP reporter expression in the eye, the brain, and the craniofacial cartilage in 10 out of 14 transgenic tadpoles at NF stage 42/43 (Figure 4I & 4J), in line with the findings in *Xenopus tropicalis.* When the reporter construct was introduced in the dorsal blastomeres at the four-cell stage, EGFP reporter expression in the eye and brain was observed in 17 out of 19 transgenic tadpoles (Figure 4K & 4L). The SED vector reporter construct containing the shared duplicated region downstream of *IRX1* (IRX1_cCRE10) also demonstrated comparable EGFP reporter expression, albeit with a much lower intensity (Figure 4M & 4N). Regarding the cCREs located in the *PRDM13* locus, the reporter construct containing the first mutational hotspot (PRDM13_cCRE3) region did not drive any EGFP reporter expression in the transgenic embryos at NF stage 42/43 (Figure 4T), while EGFP reporter expression in the eye and brain was observed for PRDM13_cCRE1 (Figure 4R & 4S) and the second mutational hotspot (PRDM13_cCRE5) (Figure 4U & 4V). This is in line with previous findings of *in situ* hybridization in *Xenopus laevis* embryos, demonstrating that *prdm13* expression is observed in progenitor cells of the developing retina, from NF stage 28 onwards. At NF stage 42/43, *prdm13* expression is detected in the amacrine cells, located in the inner part for the inner nuclear layer.^80, 81^ Moreover, for the five cCREs analyzed using these enhancer detection assays, DNase-seq profiles generated in human embryonic retinal tissue at five different developmental stages, generated by Meuleman *et al*. (2020)^46^, were analyzed in more detail. In case of IRX1_cCRE7, IRX1_cCRE10, PRDM13_cCRE1, and PRDM13_cCRE7, open chromatin is observed at/until day 103 of development, while PRDM13_cCRE1 is closed exclusively at this period. This developmental stage corresponds with the moment when retinal progenitor cells of the macula exit mitosis and differentiate towards photoreceptor fate, suggesting a functional effect of these cCREs during this period. A summary of the findings of the *Xenopus laevis* transgenesis experiments is given in Table S9.

### Novel non-coding SNVs found in the mutational hotspots of *PRDM13*

In order to expand the mutational spectrum of NCMD, twenty-three unrelated index cases with a clinical diagnosis of NCMD underwent targeted sequencing and copy number profiling of the known hotspots of the *PRDM13* and *IRX1* regions, followed by WGS for a subset of nine undiagnosed cases after targeted testing. In the *PRDM13* locus, this revealed two novel non-coding SNVs, called V15 and V16, as well as a previously reported NCMD-associated SNV (V1). Copy number analysis via qPCR did not reveal any additional variants in the two disease loci. A summary of the genetic findings can be found below and in Table S10. An overview of the pedigrees and clinical details of the families with (likely) pathogenic variants can be found in Figures S5 (family F1), S6 (family F2), and S7 (family F3), and in Table S10.

More specifically, in the index case of a Czech family (F1-III:1), a novel heterozygous SNV was found in the second mutational hotspot of *PRDM13* (V15, chr6:99599064A>G), approximately 150 bp downstream of the previously reported V10 and V11 variants (Figure 5). Subsequent segregation analysis confirmed the variant in F1-III:1 and demonstrated the presence of this variant in the two affected daughters (F1-IV:1 and F1-IV:2) and the affected paternal aunt (F1-II:3). The variant was absent in the unaffected son of F1-II:3 (F1-III:5). A second novel heterozygous SNV was identified in the index case from a Mexican family (F2-II:1). This SNV was found at the same nucleotide position as the previously reported V1 variant, albeit resulting in a different nucleotide change (V16, chr6:99593030G>C) (Figure 5). This variant, located in the first mutational hotspot of *PRDM13*, was present in three affected family members (F2-I:1, F2-III:1, and F2-III:2) and absent in three unaffected siblings (F2-II:2, F2-II:3, and F2-II:4). In the index case of another American family (F3-III:2), the known heterozygous V1 variant (chr6:99593030G>T) was identified, which was absent in two unaffected family members (F3-III:1 and F3-IV:1) and segregated in two affected family members (F3-II:1 and F3-II:3). All three reported variants are absent from gnomAD. In the twenty other families, no potential (likely) pathogenic variants were identified in the two NCMD-associated disease loci. Moreover, WGS in nine unsolved NCMD index cases did not reveal (likely) pathogenic variants in the coding regions of 290 known retinal disease genes (RetNet panel V5).

**Figure 5.**
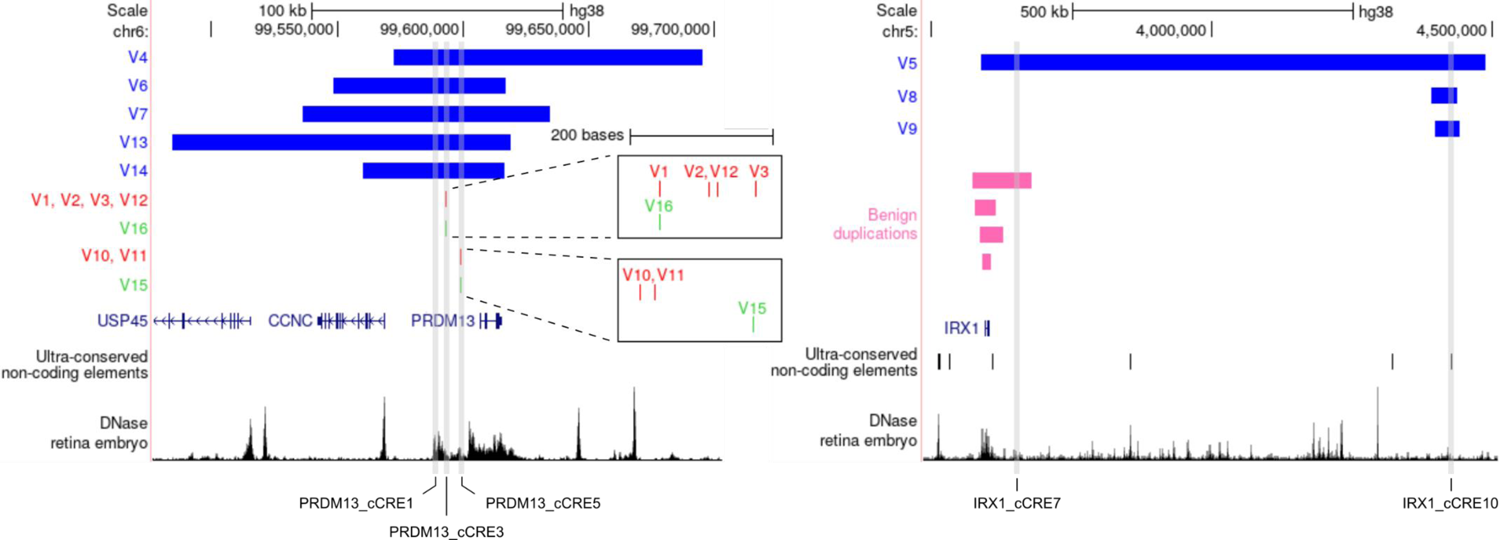
Overview of previously reported and novel (likely) pathogenic variants (V) in the *PRDM13* locus (left) and *IRX1* locus (right). Known and novel SNVs are indicated by red and green bars, respectively. The pathogenic tandem duplications are shown as blue bars, while benign duplications spanning the *IRX1* coding region, derived from DGV, are shown as pink bars. The five cCREs analyzed via *in vivo* enhancer assays in *Xenopus* are highlighted by grey vertical bars. The DNase-seq track was generated in human embryonic retinal tissue at day 103 of development. Chromosome coordinates are in hg38 annotation. DGV: Database of Genomic Variants.

### Variable *in silico* predictions of NCMD-associated SNVs

Using TRANSFAC, the effect of the eight previously reported and novel NCMD-associated non-coding SNVs on consensus TFBS was analyzed. This revealed that the novel SNV V15 is predicted to lead to the gain of an HSF1 TFBS, when compared to the wild-type sequence; while in case of the previously reported V10 variant, the A to C nucleotide change results in a predicted loss of an POU2F1 (OCT1) TFBS (Figure S8). For the six other known or novel SNVs, no effect on consensus TFBS was observed.

The regBase_REG model, predicting the regulatory potential of a variant regardless of its functional direction and pathogenicity, generated relatively high scores for all variants in both hotspots. The regBase_PAT model on the other hand, which predicts the pathogenic capacity of a variant, scores the variants in first mutational hotspot (PRDM13_cCRE3) as more likely to be pathogenic, compared to those located in the second mutational hotspot (PRDM13_cCRE5). According to GenoCanyon, which uses a cut-off of 0.5 to define functionality, the five variants located in first mutational hotspot are all functional, having a score of 1.0. This is in contrast with the three variants in the second mutational hotspot, which have low GenoCanyon scores. A similar result is obtained from the ncER prediction tool, which generates a score ranging from 0 (non-essential) to 100 (putative essential). There, the variants in the two hotspots have a score of ∼70-80 and ∼10-30, respectively. The DVAR scores, reflecting the probability of a variant being functional, are ∼0.9 for variants in first and ∼0.7 for variants in the second mutational hotspot. The PHRED-scaled FIRE scores of all variants were comparable with those from DVAR, with higher scores suggesting greater capability to regulate nearby gene expression levels. Based on the PHRED-scaled PAFA scores, only variants V1 and V16, located in the first mutational hotspot (PRDM13_cCRE3), are predicted to have a functional effect. Finally, the CDTS scores demonstrate moderate intolerance to variation for all variants (Table 2).

**Table 2.**
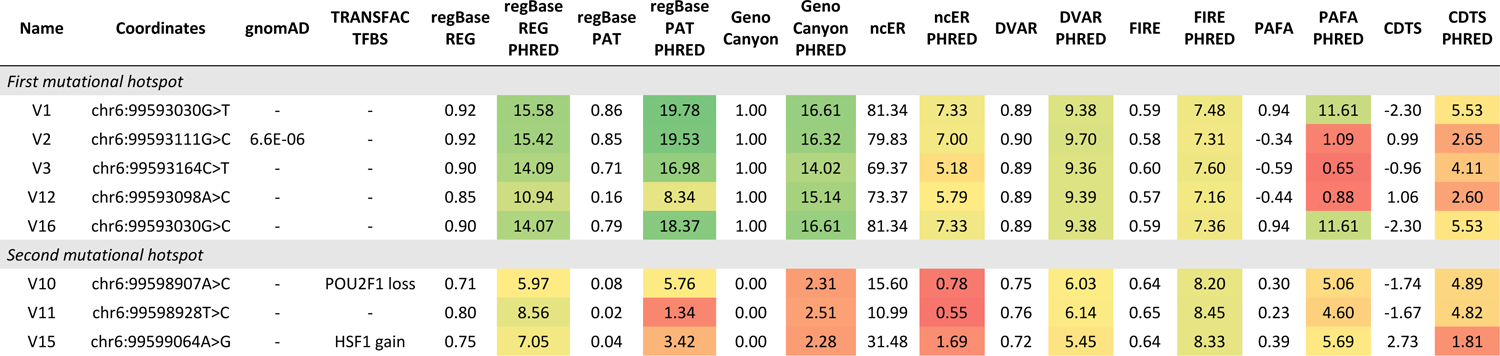
Overview of the previously reported and novel NCMD-associated SNVs (V) in the two mutational hotspots upstream of *PRDM13*. For each variant, the coordinates (hg38), nucleotide change, gnomAD v3.1.2 frequency, and TRANSFAC output are given, as well as the prediction scores and the PHRED-scaled scores 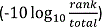 of seven *in silico* tools. The regBase_REG and regBase_PAT scores range from 0 to 1, with higher values indicating that a variant has a higher regulatory or pathogenic potential, respectively. GenoCanyon scores range from 0 to 1. A cutoff of 0.5 was used to define functionality. The ncER score ranges from 0 to 100, with higher scores suggesting that a region is more vital in terms of regulation. DVAR scores, ranging from 0 to 1, can be interpreted as the probability of a variant being functional. The FIRE score also ranges from 0 to 1, with higher scores providing stronger evidence that a variant regulates nearby gene expression levels. PAFA scores range from −1 to 1, reflecting the association of a variant with complex diseases or trait-associated SNPs. The CDTS score represents the absolute difference of the observed variation from the expected variation, with lower scores suggesting more intolerance to variation. The normalized, PHRED-scaled scores are color-coded for easy comparison (green = high, red = low).

### *In vitro* enhancer assays show a regulatory effect for NCMD-associated variants in the *PRDM13* and *IRX1* region

Next, the potential effect of the eight non-coding NCMD-associated SNVs on downstream reporter activity was assessed. Therefore, the two mutational hotspot-containing luciferase reporter vectors were subjected to site-directed mutagenesis to create eight individual variant vectors (Table S4). The luciferase assays revealed that the four known variants (V1, V2, V3, V12) in the first mutational hotspot (PRDM13_cCRE3) respectively resulted in a 2.6-, 2.8-, 1.6-, and 2.0-fold increase of reporter expression (P<0.001), relative to their wild-type vector. Similar effects were obtained for the novel variant (V16) in the same region. There, the relative luminescence was 3.2-fold higher (P<0.001) compared to the wild-type vector (Figure 6A & Table S8). For the variants located in second mutational hotspot (PRDM13_cCRE5), an opposite trend was observed. The two known (V10, V11) and one novel variant (V15) in this region respectively demonstrated a 0.6-, 0.7-, and 0.6-fold decrease of luciferase expression (P<0.001), relative to their wild-type vector (Figure 6B & Table S8).

**Figure 6.**
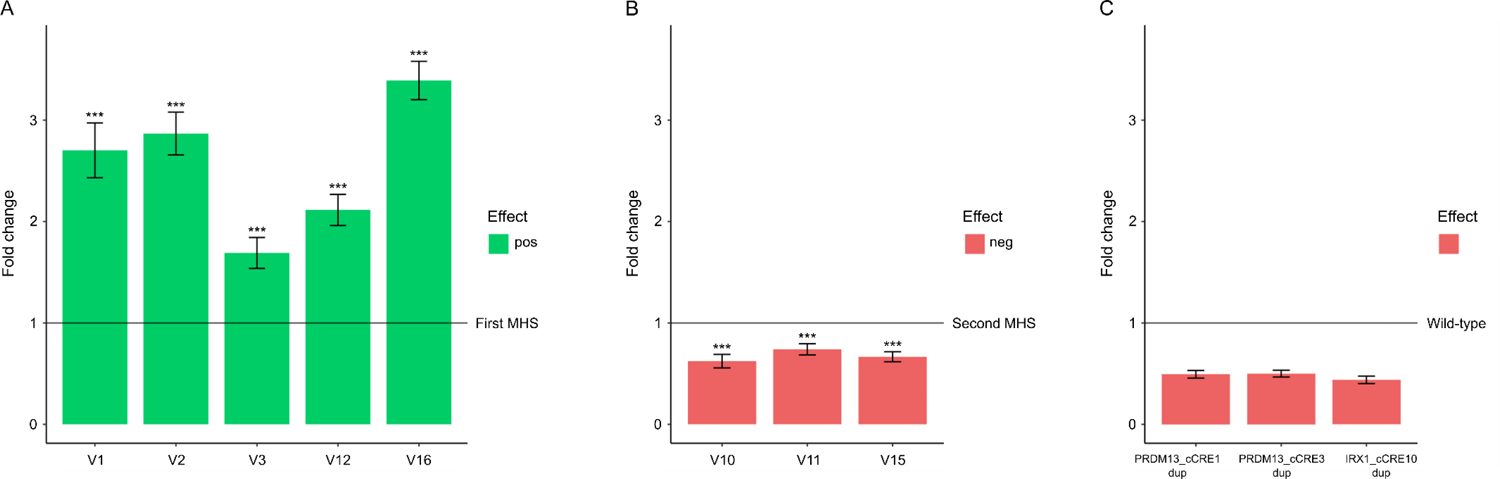
Overview of the *in vitro* mutant versus wild-type luciferase assays in ARPE-19 cells. (**A**) In the first mutational hotspot, the four known (V1-V3, V12) and one novel variant (V16) demonstrate a significant increase of luciferase reporter activity (P<0.001), relative to the wild-type vector. (**B**) In contrast, the two known (V10, V11) and one novel variant (V15) located in the second mutational hotspot, cause a significant decrease of luciferase reporter activity (P<0.001), relative to the wild-type vector. (**C**) For three non-coding regions of interest, located in the shared duplicated region of either the *PRDM13* or *IRX1* locus, the level of luciferase reporter was reduced by half (P<0.001) when the region was present as tandem duplication, relative to their respective wild-type counterpart, containing the same region of interest as a single insert. MHS: mutational hotspot.

To evaluate whether the cCREs located in previously reported NCMD-associated tandem duplications could have altered effects on reporter gene expression upon duplication, we analyzed luciferase activity of three regions located in the shared duplicated region of either the *PRDM13* or *IRX1* locus (PRDM13_cCRE1, PRDM13_cCRE3, IRX1_cCRE10). Each of these regions was cloned into a separate luciferase reporter vector (pGL4.23) as a tandem duplication, to be compared against its reciprocal vector containing the same region as single insert. The results from the luciferase assays show that in case of all three cCREs, a 0.5-fold decrease in expression levels of the luciferase reporter (P<0.001) is exhibited by the duplicated region in comparison with its single region counterpart (Figure 6C & Table S8).

### Tandem duplications in the *PRDM13* and *IRX1* loci are caused by nonhomologous end-joining or replicative-based mechanisms

To assess the underlying mechanisms that gave rise to the eight previously reported NCMD-associated tandem duplications in the *PRDM13* and *IRX1* loci, bioinformatic analyses were performed on the breakpoint regions of these duplications. We found that three out of eight tandem duplication showed overlap with repetitive elements at both breakpoints. These pairs of elements, however, had little mutual sequence identity, since they were part of different classes. Apart from repetitive elements, other sequence motifs have also been shown to predispose to DNA breakage. Therefore, we investigated the 70 bp sequence surrounding the exact breakpoints of the eight tandem duplications for the presence of 40 sequence motifs previously associated with DNA breakage. Interestingly, one particular motif (deletion hotspot consensus, TGRRKM) was present in at least one breakpoint region of seven out of eight tandem duplications, while for four of these, this motif was present in both breakpoint regions. Moreover, the inclusion of several random nucleotides at the joint point (i.e. information scar) was observed at three tandem duplications, whereas a microhomology of two or three bps was present at six tandem duplications. Based on these results, we conclude that the tandem duplications are caused either by nonhomologous end-joining (NHEJ) or by a replicative-based mechanism, such as fork stalling and template switching (FoSTeS) or microhomology-mediated break-induced replication (MMBIR). In contrast to replicative-based mechanisms, NHEJ does not require microhomology, but is generally associated with the presence of an information scar (V4, V5, V6). Microhomology in the absence of an information scar in all other tandem duplications favors the hypothesis of a replicative-based mechanism. The sites of microhomology serve as priming location to invade the second replication fork after stalling or collapse of the first replication fork. For FoSTeS and MMBIR, a single DNA break and the presence of microhomology at the breakpoints are sufficient to cause this template switching in a backward direction, which could have given rise to five out of eight tandem duplications (V7, V8, V9, V13, V14). A summary of these findings can be found in Table S11, while the presence of microhomology is visualized in Figure S9.

### During retinal development *PRDM13* is predominantly expressed in amacrine cells and *IRX1* in retinal ganglion cells

Using the snRNA-seq data from six embryonic (53, 59, 74, 78, 113, and 132 days) and three adult (25, 50, and 54 years old) human retinal samples generated by Thomas *et al.* (2022)^75^, we determined the expression pattern of *PRDM13* and *IRX1* during retinal development and in adult retina. The snRNA-seq data was first clustered into ten transcriptionally distinct clusters representing the major retinal cell types: amacrine cells, astrocytes, bipolar cells, cone photoreceptors, retinal ganglion cells, horizontal cells, microglia, Müller cells, retinal progenitor cells (RPCs), and rod photoreceptors (Figure 7A). To examine the dynamics of *PRDM13* and *IRX1* expression during retinal development, the different samples were then grouped into four time points: e50 = d53 and d59, e70 = d74 and d78, e100 = d113 and d132, adult = 3 samples. This revealed that throughout development *PRDM13* is predominantly expressed in amacrines cells with some expression observed in RPCs, horizontal, and ganglion cells (Figure 7B & Figure S10A). Compared to *PRDM13*, *IRX1* generally has a lower expression level during development, with the highest expression in retinal ganglion cells and minimal expression in RPCs (Figure 7C & Figure S10B). For both genes, the highest expression is observed at e70.

**Figure 7.**
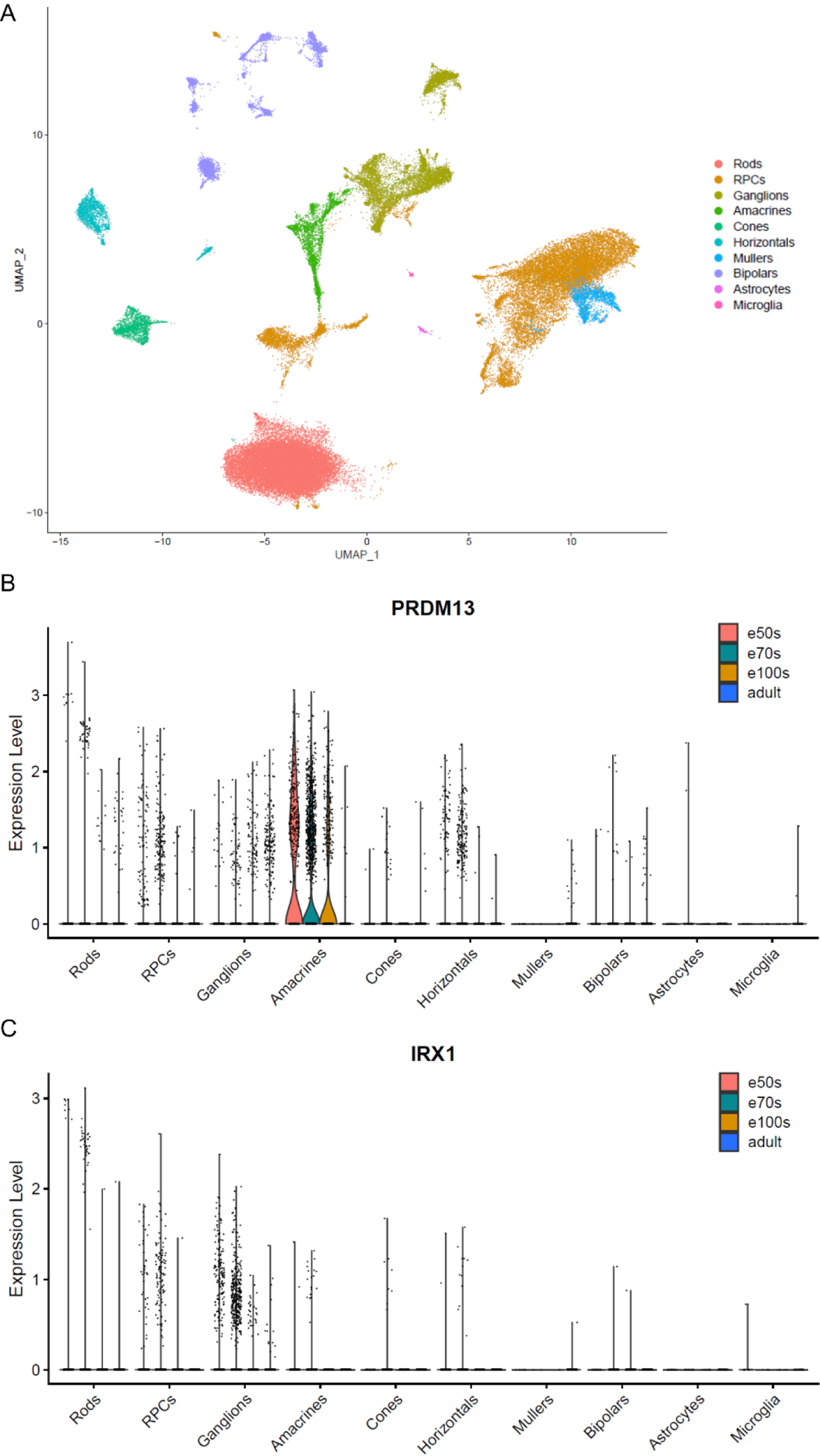
Single-cell transcriptomic analysis of developing human neural retina. In particular, data from six embryonic (e) (53, 59, 74, 78, 113, and 132 days) and three adult (25, 50, and 54 years old) human retinal samples are included. (**A**) UMAP plot of 60,014 human neural retinal cells from all samples, colored based on the ten transcriptionally distinct clusters represented in the key. (**B-C**) The different retinal samples were grouped into four time points: e50 = d53 and d59, e70 = d74 and d78, e100 = d113 and d132, adult = 3 samples. Violin plots illustrate respective *PRDM13* and *IRX1* expression in the different retinal cell types for each of these time points. RPCs: retinal precursor cells.

## Discussion

In the last decade, it has become evident that non-coding variants contribute significantly to the molecular pathogenesis of IRDs. Although an increasing number of pathogenic variants affecting *cis*-acting splicing have been identified,^19–21^ causal variants located in non-coding *cis*-acting regulatory elements are more scarce.^82–84^ The identification and interpretation of the latter, however, remains challenging as the exact mechanisms of the elements they are located in, the genes they act on, and the effect of the variation remains largely unknown. Here, we have focused on NCMD, a developmental macular disorder caused by non-coding SNVs and tandem duplications assumed to affect *PRDM13* (MCDR1 locus) and *IRX1* (MCDR3 locus) expression.

The establishment of RegRet, a genome-wide multi-omics retinal database, advances genome annotation for the human retina. It not only laid the foundation for the locus-specific research of the two genomic regions implicated in NCMD, but also represents a universal framework for studies of other loci implicated in rare IRDs as well as complex retinal diseases.

By integrating chromosome conformation capture (UMI-4C) profiles for the *PRDM13* and *IRX1* promoter, generated on adult human retinal tissue, with multiple genome-wide retinal datasets on DNA accessibility, epigenetic histone marks, transcription factor binding, and gene expression, we were able to both characterize candidate *cis*-regulatory elements (cCREs) and determine which of these physically interact with the two promoters of interest, revealing eight and ten cCRE for the *PRDM13* and *IRX1* gene, respectively. Although *in vitro* luciferase assays did not demonstrate regulatory activity for all of these regions, *in vivo* experiments in *Xenopus* embryos could confirm eye- and brain-specific activity for two cCREs in both the *PRDM13* (PRDM13_cCRE1, PRDM13_cCRE5) and the *IRX1* locus (IRX1_cCRE7, IRX1_cCRE10). Moreover, the interaction of PRDM13_cCRE1 with the *PRDM13* promoter was confirmed by a reverse UMI-4C experiment, on top of appropriate DNA accessibility and ChIP-seq data, putting forward this region as a novel enhancer of *PRDM13*. Using single-cell transcriptional mining, we showed that *PRDM13* is expressed in the fetal retina, with the highest values in the amacrine cells between day 74 and 78 of embryonic development, while little to no expression is observed in adult retinal neurons. Since we performed the UMI-4C experiments on adult human retinas, this could explain why the first mutational hotspot (PRDM13_cCRE3), which is expected to have a role during retinal development, did not show an interaction with the *PRDM13* promoter. Indeed, using the activity-by-contact (ABC) method based on Hi-C data from human embryonic stem cells, Green *et al.* (2021) demonstrated that both hotspots encompass cCREs.^85^ Interestingly, 70% of the macula-specific CREs that were identified in this study correspond with the eight candidate CREs that we identified in the *PRDM13* locus (Figure S11). It should also be noted that the quality of UMI-4C profiles, reflected by the number of UMIs, is dependent on both the genomic region the viewpoint is located in and the efficiency of the nested PCR reactions. The same holds true for the reverse UMI-4C experiment on adult retinal tissue using the shared duplicated region, located ∼800kb downstream from the *IRX1* promoter, as a bait. The obtained profile hinted at an interaction between this region and the *IRX1* promoter, while the promoter-based UMI-4C experiment did not. Similar to *PRDM13*, analysis of scRNA-seq data illustrated absence of *IRX1* expression in adult retinal tissue, while highest expression was shown between day 74 and 78 in retinal ganglion cells.

Genomic profiling of twenty-three index cases with a clinical diagnosis of NCMD revealed three non-coding SNVs upstream of the *PRDM13* gene, of which two are novel (V15, V16) and one was previously reported (V1). Variant V15 (chr6:99599064A>G), located in the second mutational hotspot of the *PRDM13* region (PRDM13_cCRE5), and segregating in three affected family members, is absent from gnomAD and has varying predictions of functionality and pathogenicity. This could be explained by the fact that most prediction tools are based on supervised modeling, and therefore heavily rely on the composition and annotation of the training dataset used, which might not sufficiently represent the type of variants under investigation. However, the nucleotide change of variant V15 causes a predicted gain of an HSF1 TFBS. For the V10 variant, located in the same hotspot, a similar effect was observed, albeit resulting in the loss of a POU2F1 (OCT1) TFBS. Interestingly, Pou2f1 has been shown to play a role in the regulation of cone photoreceptor production in mouse retina, while Hsf1 activity has been demonstrated in rat retina.^86, 87^ The second novel variant, variant V16 (chr6:99593030G>C), is located in the first mutational hotspot of the *PRDM13* region (PRDM13_cCRE3) and constitutes a different nucleotide change at the same nucleotide position as the previously reported V1 variant.^23^ For these and the three other variants in the first hotspot, *in silico* predictions support a variant effect.

Although IRDs in general have a diagnostic success rate of approximately 60%, causative variants were detected in only 13% of the NCMD cohort tested here, which could be explained by different factors. First of all, most IRD studies commonly rely on exome sequencing to identify coding variants with a high impact. Here, the search was oriented toward non-coding pathogenic variants, which is complicated by incomplete knowledge of the location and function of all regulatory elements. Furthermore, (non-coding) variants affecting other known or novel disease genes involved in retinal and/or macular development, may also be causative for NCMD. Next, NCMD has great phenotypic variability and its clinical presentation may be mimicked by phenocopies. Examples thereof are toxoplasmosis or subtypes of inherited macular disease (e.g. *BEST1*-, *PRPH2*-associated maculopathy).^36, 89^ Finally, more complex structural variants in the *PRDM13* or *IRX1* locus, affecting the regulation of target gene expression, may be missed due to technical limitations of the qPCR-based screening and short-read WGS.

When comparing both the novel and previously reported SNVs against their wild-type sequences using luciferase assays in ARPE-19 cells, we showed that all SNVs exert an effect on reporter gene expression. In case of the first hotspot (PRDM13_cCRE3), the five variants had an increased activity, with the novel V16 variant having the strongest effect, whereas the three variants in the second hotspot (PRDM13_cCRE5) all demonstrated a decreased activity. These opposite effects on reporter gene expression can be explained by the fact that the cCREs the variants are located in were tested outside of their genomic context. Their endogenous functions are expected to be more complicated than what was observed in these experiments, with combinations of them working together to fine-tune *PRDM13* gene expression. Moreover, the minimal promoter used in the pGL4.23 vector may respond differently to the cCREs compared to the *PRDM13* promoter, and transcription factors acting through them in the ARPE-19 cells might not be the same. Nonetheless, we have demonstrated that the NCMD-associated SNVs significantly affect the activity of these regulatory elements.

Both mutational hotspots have been identified as cCREs active at day 103 of embryonic development, i.e. the moment when retinal progenitor cells of the macula exit mitosis and differentiate towards photoreceptor fate,^85, 90^ and the luciferase assays show an effect of the SNVs they contain on reporter gene expression. Therefore, disruption of *PRDM13* expression during a critical step of maculogenesis can be assumed. Indeed, *Prdm13* knockout in mice and overexpression of the *PRDM13* orthologue in *Drosophila* (CG13296) presented with abnormal retinal and eye development, respectively.^34, 38^ Apart from non-coding SNVs, five NCMD-associated tandem duplications have also been identified in the *PRDM13* locus. These have a ∼44kb shared duplicated region that comprises four cCREs we identified and both mutational hotspots. Based on luciferase assays, we demonstrated that the duplication of two of these regions (PRDM13_cCRE1, PRDM13_cCRE3) also affects reporter gene expression. In addition, this region contains a CTCF binding site, indicative of a sub-TAD boundary, as determined using the RegRet database. A duplication thereof could thus lead to the generation of a neo-TAD, associated with gain-of-function effects of certain CREs, resulting in altered target gene expression.^14, 91^

Since only one of the three previously reported NCMD-associated tandem duplications in the *IRX1* locus and at least four benign duplications from DGV span the coding region of the *IRX1* gene, we can assume that the disease-causing mechanisms associated with these duplications do not result from impaired *IRX1* dosage itself, but rather have a regulatory basis. In the ∼39kb shared duplicated region downstream of *IRX1*, we identified a UCNE which, based on DNase-seq data from the developing retina, is active between day 74 and 103 of development. Similar to the observations in the hotspots of the *PRDM13* locus, this period corresponds to an important moment in human maculogenesis, characterized by the proliferation and differentiation of retinal progenitor cells.^90^ *In vivo Xenopus* experiments demonstrated eye-specific and developmental activity of this UCNE-containing region (IRX1_cCRE10), and the duplication was shown to have an effect on luciferase reporter expression *in vitro*.

By single-cell RNA sequencing of different macular subregions, Voigt *et al.* (2021) demonstrated region-specific gene expression and alternative splicing.^92^ Although this unique character of the human macula and its development are not fully represented by *in vitro* studies in postnatal cells and *in vivo* assays in a model organism lacking a macula, we have provided evidence that NCMD-associated variants can affect *PRDM13* and *IRX1* expression due to nucleotide changes in regulatory sequences, gene duplication, the duplication of one or more CREs, or the disturbance of the gene regulatory landscape. Based on these findings, we put forward a hypothesis on how transcriptional changes in two distinct loci can result in the same macular phenotype. In particular, by transcriptome profiling based on RNA-seq data from thirteen human fetal retina samples spanning different developmental stages, Hoshino *et al.* (2017) revealed three key epochs in the transcriptional dynamics of human retina between day 52 and 150.^93^ Our analysis of single-cell RNA-seq data of embryonic retina illustrated that both *PRDM13* and *IRX1* demonstrate their highest expression at the beginning of epoch two (day 67 to 80), corresponding to the moment when amacrine cells start to emerge and begin to synapse with retinal ganglion cells.^93^ We therefore hypothesize that altered *PRDM13* or *IRX1* expression impairs macula-specific synaptic interactions between amacrine and ganglion cells during retinogenesis. In future studies, transcriptome and chromatin interaction studies on patient-derived retinal organoids may be useful to evaluate the effect of SNVs and tandem duplications on gene expression, genome organization, and cellular differentiation and function *in vivo*.

In conclusion, we have provided an integrated retinal multi-omics database, which advances the annotation of the genome in human retina and represents a universal framework for the investigation of disease-associated loci implicated in rare as well as complex retinal diseases. By integrated multi-omics profiling of human retina, and by an in-depth *in silico*, *in vitro* and *in vivo* assessment of cCREs and variants therein, we have gained insight into the *cis*-regulatory mechanisms and genetic architecture of NCMD. Overall, this supports the hypothesis that NCMD is a retinal enhanceropathy, which is unique in the wider group of IRDs. With a broader implementation of WGS in rare disease research, an emphasis shift to non-coding regions such as CREs as targets of mutations can be expected. Finally, our study is exemplar for how expanding knowledge of disease-causing mechanisms and phenotype-driven genomic and functional data profiling of non-coding regions, advance our ability to fully interpret variants in non-coding regions in rare diseases. Providing a definitive genetic diagnosis in more individuals with rare diseases is imperative for the design of efficient disease-specific genetic testing, genetic counseling, and ultimately for therapeutic decisions.

## Data availability

All unique data and materials that support the findings of this study are readily available from the corresponding author upon request. The genome-wide retinal multi-omics database RegRet, composed during this study, can be accessed via the University of California, Santa Cruz (UCSC) Genome Browser: http://genome.ucsc.edu/s/stvdsomp/RegRet. The in-house RetNet panel v5 can be accessed at https://www.cmgg.be/assets/bestanden/Genpanel-RETNET-v5.pdf.

## Acknowledgements

This work was supported by grants from Ghent University Special Research Fund (BOF20/GOA/023) (EDB, KV, BPL); Ghent University Hospital Innovation Fund NucleUZ (EDB); JED Foundation (EDB); H2020 Marie Sklodowska-Curie Innovative Training Networks (ITN) StarT (grant No. 813490) (EDB, KV, JLGS, JJT, JRM); EJP RD Solve-RET EJPRD19-234 (EDB, PL, BK, CR, JLGS, JJT, JRM), SNSF grant # 204285 (CR), Foundation Fighting Blindness in Columbia, MD (Grant #: BR-GE-1216-0715-CSH). SVS (1145719N) is PhD fellow of the Research Foundation-Flanders (FWO), EDB (1802220N) and BPL (1803816N) are FWO Senior Clinical Investigators; MBS, VLS and ADR are Early Starting Researcher of ITN StarT (grant No. 813490). BK, BPL, CJFB, EDB, PL and VV are members of ERN-EYE (Framework Partnership Agreement No 739534-ERN-EYE). An unrestricted grant from The Molecular Insight Research Foundation (KWS).

## Author contributions

SVS: Conception and project design, acquisition of data, analysis and interpretation of data, drafting and revising the manuscript.

KWS: Acquisition of data, analysis and interpretation of data, drafting and revising the manuscript.

MBC: Project design, acquisition of data, analysis and interpretation of data, drafting and revising the manuscript.

VLS: Project design, acquisition of data, analysis and interpretation of data, drafting and revising the manuscript.

ED: Project design, acquisition of data, analysis and interpretation of data, revising the manuscript.

FSS: Acquisition of data, analysis and interpretation of data, revising the manuscript.

SA: Acquisition of data, analysis and interpretation of data, revising the manuscript.

TVDS: Acquisition of data, analysis and interpretation of data, revising the manuscript.

ADR: Acquisition of data, analysis and interpretation of data, revising the manuscript.

TR: Acquisition of data, revising the manuscript.

MVH: Acquisition of data, revising the manuscript.

SV: Analysis and interpretation of data, revising the manuscript.

IB: Acquisition of data, revising the manuscript.

AAB: Acquisition of data, revising the manuscript.

CJFB: Acquisition of data, revising the manuscript.

JDZ: Acquisition of data, revising the manuscript.

CFI: Acquisition of data, revising the manuscript.

BK: Acquisition of data, revising the manuscript

VV: Acquisition of data, revising the manuscript

BPL: Acquisition of data, revising the manuscript.

CR: Acquisition of data, revising the manuscript.

JvdE: Acquisition of data, revising the manuscript.

MJvS: Acquisition of data, revising the manuscript.

VV: Acquisition of data, revising the manuscript.

JLGS†: Project design, acquisition of data, analysis and interpretation of data.

JJT: Project design, acquisition of data, analysis and interpretation of data, revising the manuscript.

JRMM: Project design, acquisition of data, analysis and interpretation of data, revising the manuscript.

PL: Acquisition of data, analysis and interpretation of data, drafting and revising the manuscript.

KV: Project design, acquisition of data, analysis and interpretation of data, revising the manuscript.

EDB: Conception and project supervision, acquisition of data, analysis and interpretation of data, drafting and revising the manuscript.

## Supplementary data

**Table S1.**
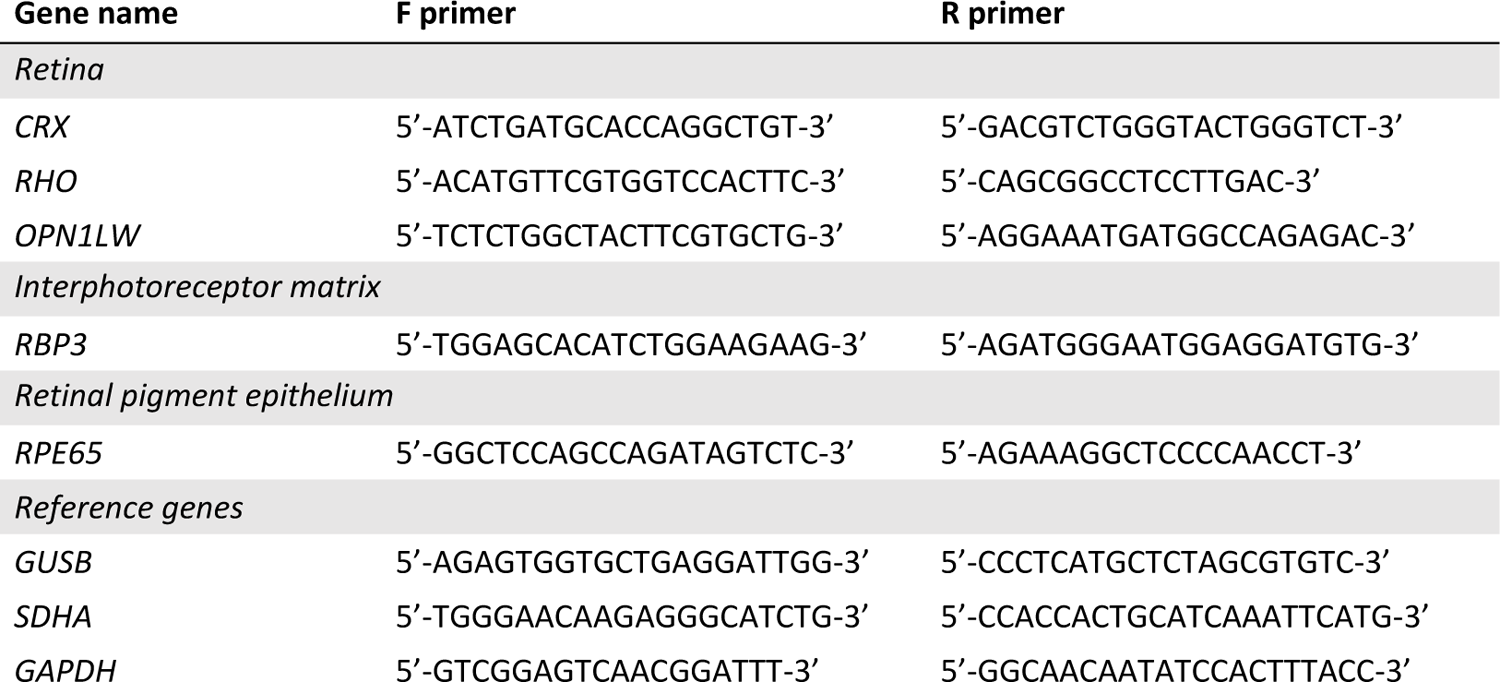
Expression primers. Overview of the qPCR primers used for assessing the purity of the retinal tissue on which UMI-4C was performed.

**Table S2.**
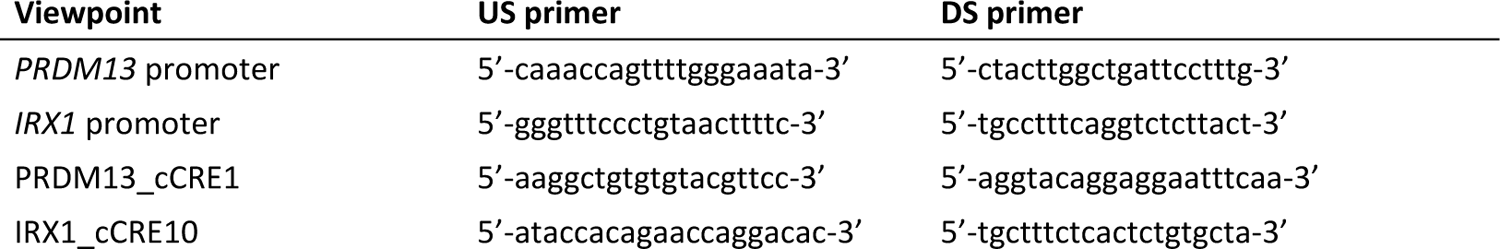
UMI-4C primers. Overview of the US (first PCR reaction) and DS (second PCR reaction) primers used for generating the UMI-4C libraries.

**Table S3.**
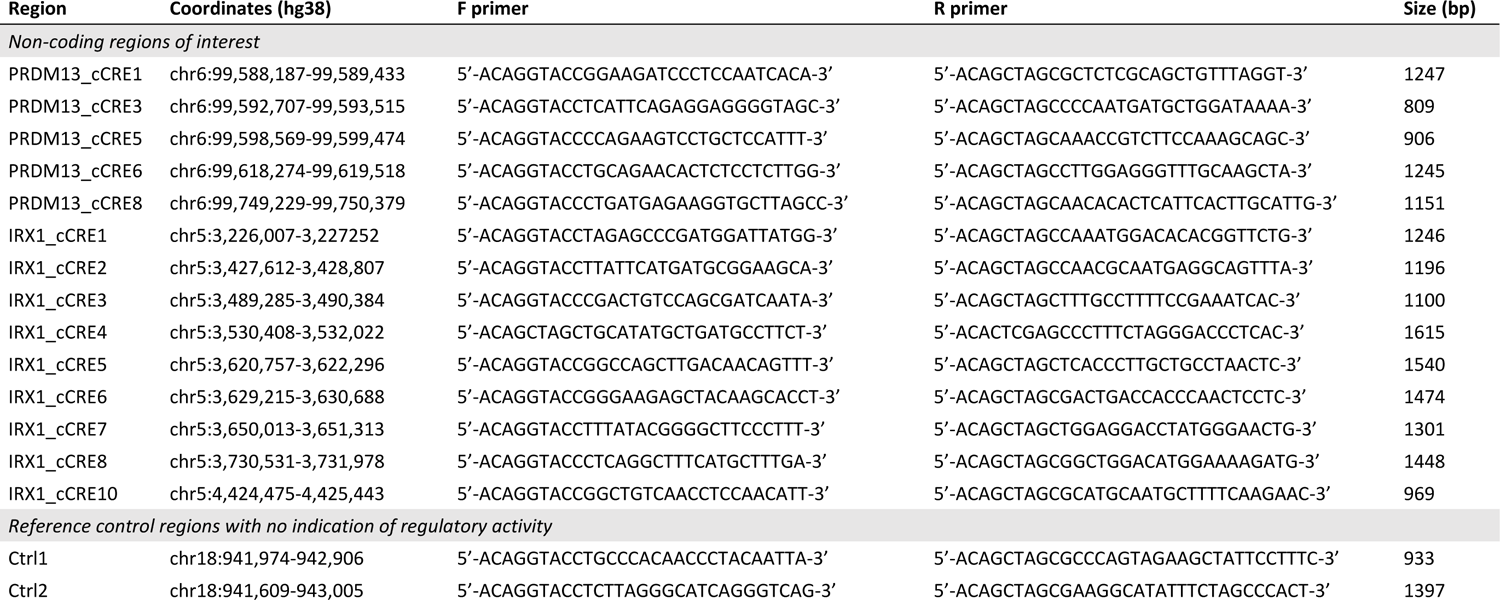
*In vitro* reporter assay constructs. Overview of the genomic regions that were cloned into pGL4.23 luciferase reporter vectors, together with their respective PCR primers.

**Table S4.**
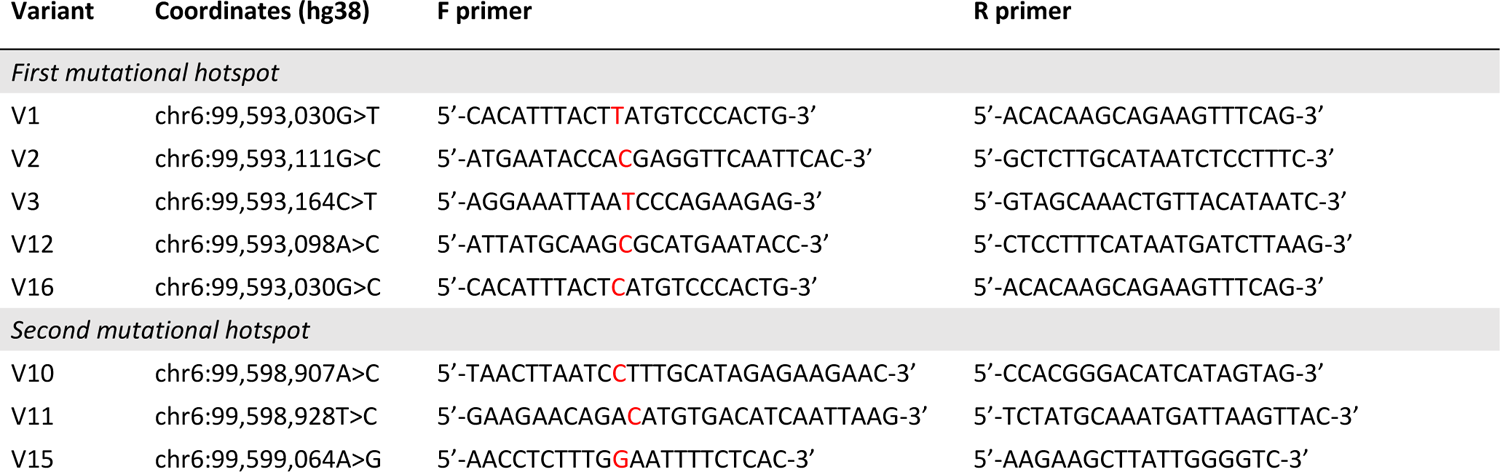
Mutagenesis primers. Overview of the mutagenesis primers used for introducing the different SNVs into pGL4.23 luciferase reporter vectors containing the respective region of interest.

**Table S5.**
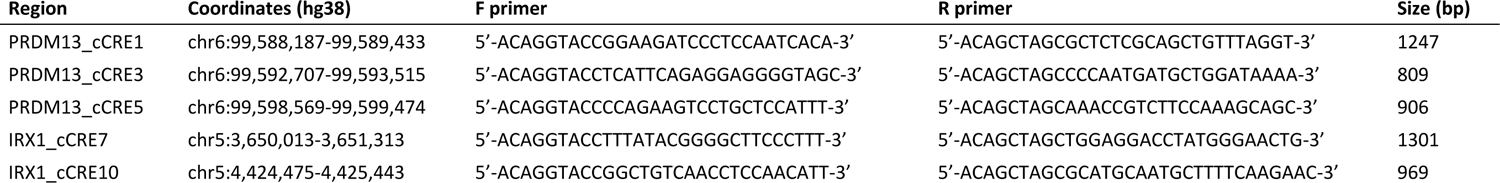
*In vivo* reporter assay constructs. Overview of the non-coding regions of interest that were cloned into SED reporter vectors, together with their respective PCR primers.

**Table S6.**
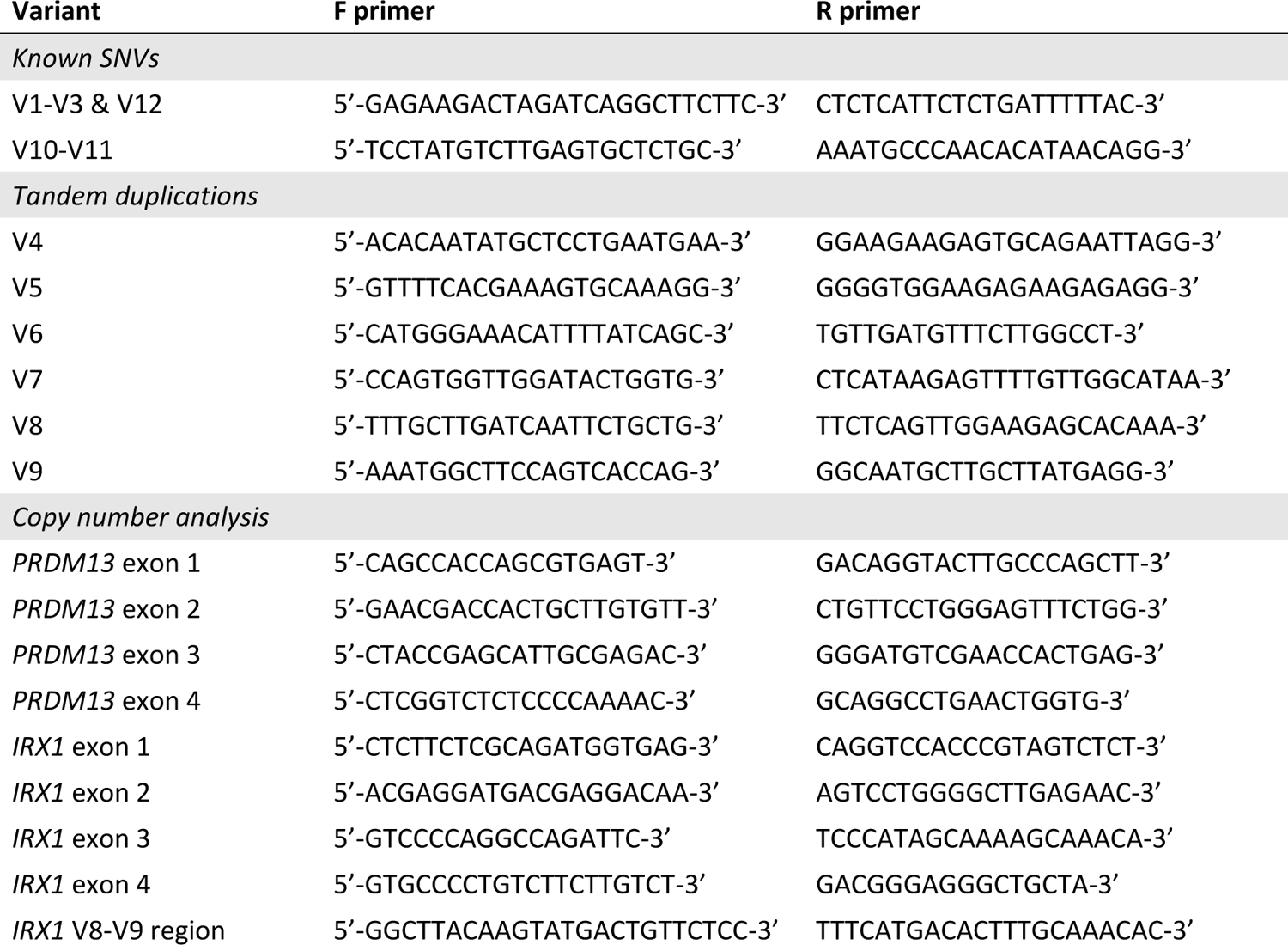
Targeted genetic testing primers. Overview of the PCR primers used for targeted testing the known *PRDM13*-associated SNVs and tandem duplications (*PRDM13* and *IRX1*), and the qPCR primers used for performing the copy number analysis in the two NCMD-associated disease loci *PRDM13* and *IRX1*.

**Table S7.** UMI-4C read count. Number of raw, mapped, and unique reads from UMI-4C sequencing of the different viewpoints.

| Viewpoint | Sample | Raw | Mapped | UMIs |
| --- | --- | --- | --- | --- |
| *PRDM13* promoter | Donor 1 | 8,134,307 | 687,131 | 13,276 |
|  | Donor 2 | 7,330,735 | 290,656 | 20,842 |
| *IRX1* promoter | Donor 1 | 8,498,021 | 1,894,101 | 9,569 |
|  | Donor 2 | 4,890,848 | 1,555,296 | 15,694 |
| PRDM13_cCRE1 | Donor 1 | 4,445,807 | 967,751 | 7,150 |
|  | Donor 2 | 6,474,247 | 733,233 | 11,331 |
| IRX1_cCRE10 | Donor 1 | 9,074,572 | 3,277,923 | 13,943 |
|  | Donor 2 | 7,542,129 | 2,704,943 | 18,199 |

**Table S8.** Results from the luciferase assays. For the known and novel SNVs, the three artificial tandem duplications, and the set of fourteen non-coding regions of interest, the fold change of the luciferase reporter level relative to the level of the reference control luciferase vector is given. For the variants, the P-values were obtained by likelihood ratio tests of the linear mixed effects model with the fixed effect against a null model without this effect. cCRE: candidate CRE, MHS: mutational hotspot, pos: positive, NS: not significant, neg: negative.

| Insert | Effect | Fold change | P-value |
| --- | --- | --- | --- |
| *Non-coding region of interest* |  |  |  |
| PRDM13_cCRE1 | NS | 1.02 ± 0.07 | 8.1E-01 |
| PRDM13_cCRE3 | pos | 1.90 ± 0.08 | 6.2E-09 |
| PRDM13_cCRE5 | NS | 0.88 ± 0.07 | 8.0E-02 |
| PRDM13_cCRE6 | pos | 2.57 ± 0.08 | 6.7E-13 |
| PRDM13_cCRE8 | NS | 0.98 ± 0.08 | 8.2E-01 |
| IRX1_cCRE1 | pos | 3.04 ± 0.10 | 6.2E-14 |
| IRX1_cCRE2 | pos | 2.07 ± 0.08 | 1.6E-10 |
| IRX1_cCRE3 | pos | 1.38 ± 0.08 | 2.1E-04 |
| IRX1_cCRE4 | neg | 0.62 ± 0.06 | 2.1E-05 |
| IRX1_cCRE5 | neg | 0.49 ± 0.07 | 1.1E-06 |
| IRX1_cCRE6 | NS | 0.96 ± 0.08 | 6.4E-01 |
| IRX1_cCRE7 | pos | 6.82 ± 0.32 | 1.2E-12 |
| IRX1_cCRE8 | pos | 2.44 ± 0.15 | 2.2E-08 |
| IRX1_cCRE10 | NS | 1.12 ± 0.06 | 6.1E-02 |
| *SNVs in first mutational hotspot* |  |  |  |
| V1 | pos | 2.58 ± 0.27 | 2.4E-05 |
| V2 | pos | 2.75 ± 0.21 | 2.6E-07 |
| V3 | pos | 1.62 ± 0.15 | 5.1E-04 |
| V12 | pos | 2.03 ± 0.13 | 6.0E-07 |
| V16 | pos | 3.24 ± 0.19 | 1.9E-09 |
| *SNVs in second mutational hotspot* |  |  |  |
| V10 | neg | 0.59 ± 0.07 | 1.2E-05 |
| V11 | neg | 0.70 ± 0.07 | 3.5E-04 |
| V15 | neg | 0.64 ± 0.07 | 8.1E-05 |
| *Tandem duplications* |  |  |  |
| PRDM13_cCRE1_dup | neg | 0.52 ± 0.03 | 2.1E-12 |
| PRDM13_cCRE3_dup | neg | 0.53 ± 0.03 | 5.7E-12 |
| IRX1_cCRE10_dup | neg | 0.46 ± 0.04 | 6.6E-11 |

**Table S9.** Summary of the results from the *in vivo* transgenic enhancer assays in *Xenopus*. In *Xenopus tropicalis* and albino *Xenopus laevis*, *in vivo* transgenic enhancer assays were performed using a fluorescent enhancer detection vector and transgenesis via I-*Sce*I meganuclease. For each of the five assayed cCREs of interest, it is indicated where EGFP reporter expression was observed at five different developmental stages. NF: Nieuwkoop and Faber, NC: neural crest, NE: no expression, NDA: no data available.

| Region | Species | NF stage 15 | NF stage 20 | NF stage 42/43 | NF stage 45 | NF stage 55 |
| --- | --- | --- | --- | --- | --- | --- |
| IRX1_cCRE7 | *X. tropicalis* | Neural plate | Neural tube | NC derivatives, eye | NC derivatives, eye | Eye |
|  | *X. laevis* | NE | NE | NC derivatives, eye, brain | NDA | NDA |
| IRX1_cCRE10 | *X. tropicalis* | NE | NE | NDA | NDA | Eye |
|  | *X. laevis* | NE | NE | Eye, brain | NDA | NDA |
| PRDM13_cCRE1 | *X. tropicalis* | NE | NE | Eye | NDA | NDA |
|  | *X. laevis* | NE | NE | Eye, brain | NDA | NDA |
| PRDM13_cCRE3 | *X. tropicalis* | NDA | NDA | NDA | NDA | NDA |
|  | *X. laevis* | NE | NE | NE | NE | NE |
| PRDM13_cCRE5 | *X. tropicalis* | NE | NE | Eye | Eye | NDA |
|  | *X. laevis* | NE | NE | Eye, brain | NDA | NDA |

**Table S10.** Summary of genetic and clinical findings of the families with (likely) pathogenic variants. BCVA: best-corrected visual acuity, OCT: optical coherence tomography, RE: right eye, LE: left eye, ERG: electroretinogram, EOG: electrooculogram, unk: unknown.

| Family, pedigree number | Variant | Age/sex | BCVA | Fundus imaging | OCT | Other findings | NCMD grade |
| --- | --- | --- | --- | --- | --- | --- | --- |
| F1, III:1 | V15 | 36/F | RE: 20/20<br>LE: 20/20 | Yellowish-white lesions in the macula, encircled by hypopigmented halos and surrounded by hyperpigmentation | Focal RPE damage and focal neurosensory detachment | Normal color vision | 2 |
| F1, IV:1 | V15 | 18/F | RE: 20/20<br>LE: 20/80 | Circular plaques of coalescing yellowish-white masses in the macula and gross pigment deposits | Focal RPE damage, right focal neurosensory detachment and left submacular scarring | Normal color vision | 2 |
| F1, IV:2 | V15 | 12/F | RE: 20/20<br>LE: 20/20 | Mild RPE abnormalities with small macular drusen. | Mild RPE abnormalities with small macular drusen | Normal color vision | 2 |
| F1, II:3 | V15 | 73/F | RE: 20/50<br>LE: 20/160 | Chorioretinal macular atrophy with fibrotic changes more pronounced in the LE | Chorioretinal macular atrophy with fibrotic changes more pronounced in the LE, and vitreofoveal adhesion | - | 3 |
| F2, II:1 | V16 | 41/F | RE: 20/30<br>LE: 20/40 | Symmetrical elevated submacular yellow vitelliform-like lesions | Vitelliform lesions with central fibrosis | - | 2 |
| F2, III:1 | V16 | 15/F | RE: 20/50<br>LE: 20/63 | Symmetrical coloboma-like macular malformations | - | Flat/abnormal EOG, normal full-field ERG | 3 |
| F2, III:2 | V16 | 12/M | RE: 20/70<br>LE: 20/80 | Symmetrical coloboma-like macular malformations | - |  | 3 |
| F3, III:2 | V1 | 37/F | RE: 20/200<br>LE: 20/200 | Severe bilateral lesions of the RPE affecting the entire macular area | Excavated lesion with a temporal shelving edge and subretinal fibrosis | - | 3 |
| F3, II:1 | V1 | unk/F | RE: 20/25<br>LE: 20/30 | Yellow specks in the central macula of both eyes | - | - | 1 |
| F3, II:3 | V1 | unk/M | RE: 20/30<br>LE: 20/25 | Yellow specks in the central macula of both eyes | - | - | 1 |

**Table S11.** Summary of the bioinformatics analyses of the eight previously reported NCMD-associated tandem duplications. For the breakpoint regions of each duplication, the degree of microhomology, the presence of repetitive elements, and the presence of sequence motifs, based on 40 previously described motifs (Fuzznuc) are indicated.

|  | V4 | V5 | V6 | V7 | V8 | V9 | V13 | V14 |
| --- | --- | --- | --- | --- | --- | --- | --- | --- |
| Chr | 6 | 5 | 6 | 6 | 5 | 5 | 6 | 6 |
| Start (hg38) | 99,572,329 | 3,587,787 | 99,548,350 | 99,536,433 | 4,391,264 | 4,396,814 | 99,484,588 | 99,560,265 |
| End (hg38) | 99,695,430 | 4,485,914 | 99,617,261 | 99,634,822 | 4,436,422 | 4,440,329 | 99,619,234 | 99,616,492 |
| Inserted bases (bp) | 22 | 15 | 5 | 0 | 0 | 0 | 0 | 0 |
| Microhomology (bp) | 0 | 2 | 0 | 2 | 2 | 2 | 3 | 3 |
| *5' breakpoint region* |  |  |  |  |  |  |  |  |
| Repetitive element | AluSx1 | - | MER117 | AluJb | - | - | AluJr | - |
| Number of sequence motifs | 2 | 6 | 5 | 7 | 4 | 2 | 1 | 2 |
| *3' breakpoint region* |  |  |  |  |  |  |  |  |
| Repetitive element | L1M1 | - | AluJo | L2a | L1M1 | L1MC3 | - | - |
| Number of sequence motifs | 3 | 1 | 4 | 5 | 3 | 3 | 1 | 2 |
| *Conclusion* |  |  |  |  |  |  |  |  |
| Potential molecular mechanism | NHEJ | NHEJ | NHEJ | Replicative | Replicative | Replicative | Replicative | Replicative |

---

**Figure S1.**
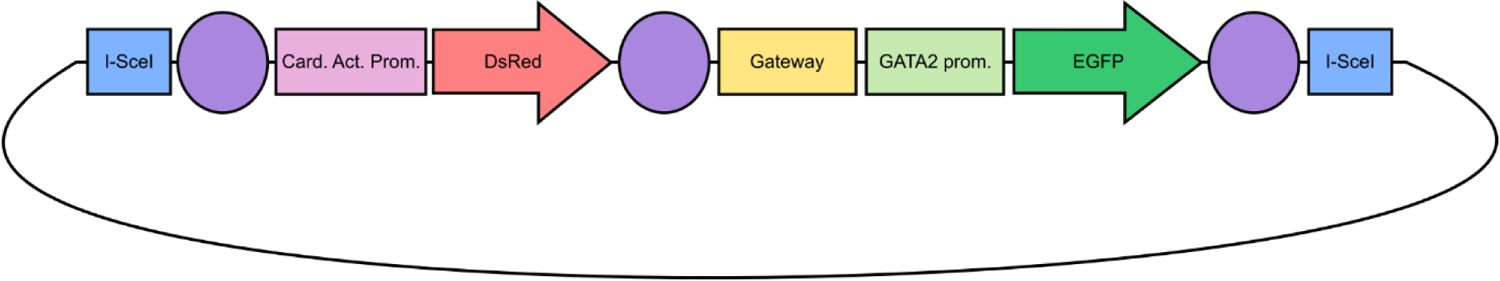
Schematic representation of the SED vector. The transgenesis internal control cassette is composed of the Cardiac Actin promoter (pink), driving strong expression in the somites, and the DsRed fluorescent protein (red), serving as a control for transgenesis efficiency *in vivo* in the F0 and the F1 embryos. The enhancer detection cassette contains a Gateway entry site (yellow), the *gata2* minimal promoter (light green), and the enhanced green fluorescent protein (EGFP) reporter gene (dark green). EGFP reporter expression can be observed during early development under a fluorescent microscope. Both cassettes are flanked by insulator sequences (purple) to protect them from position effects, and together they are flanked by I-*Sce*I meganuclease recognition sites (blue).

**Figure S2.**
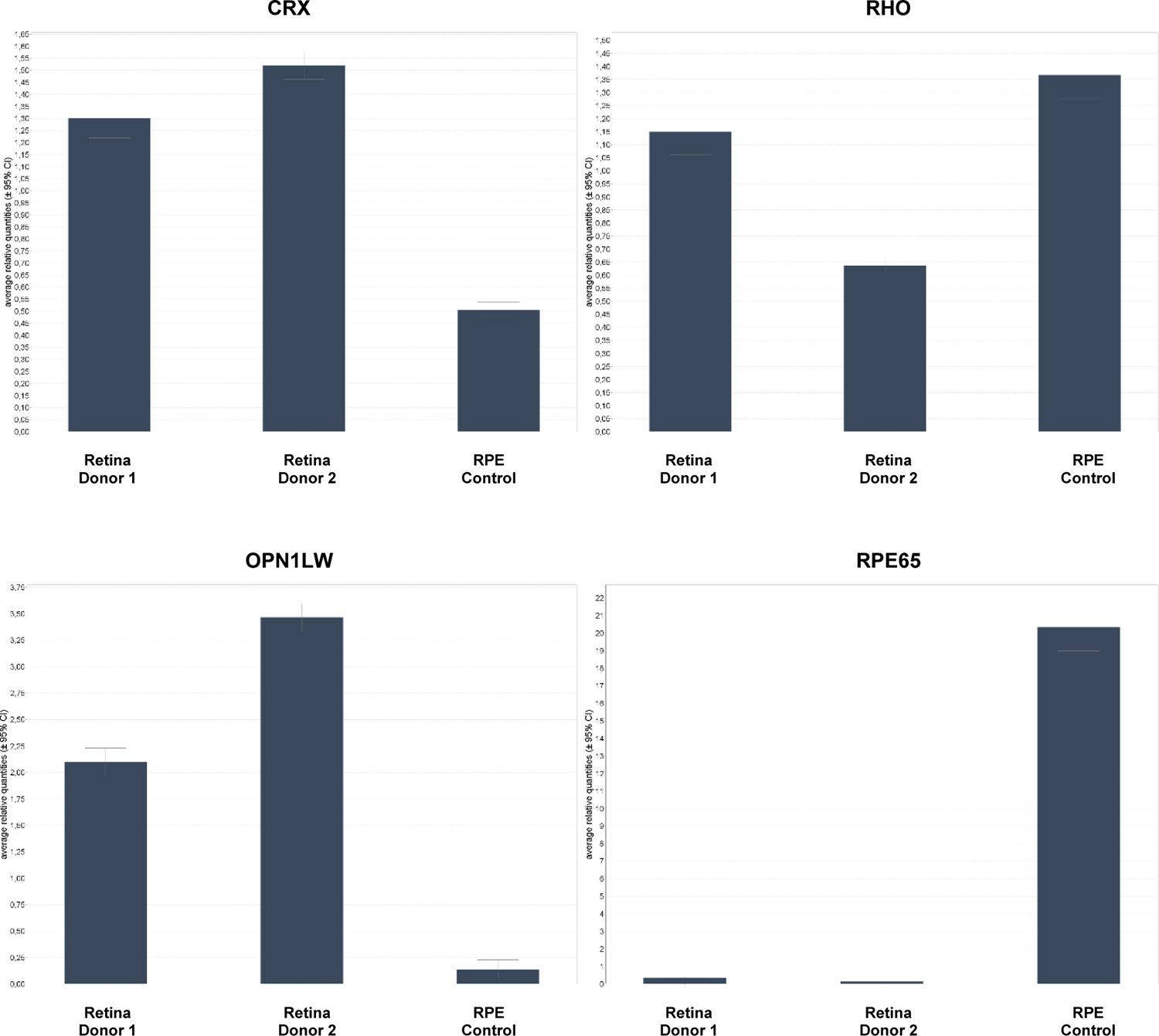
Post-dissection expression analysis of human donor retina. Output from the qPCR expression analysis of the two dissected retinal samples, compared against a control sample of RPE tissue. The retinal genes *CRX*, *RHO*, and *OPN1LW* are highly expressed in the retinal tissues, while expression of the RPE-specific *RPE65* gene is minimal relative to the RPE sample.

**Figure S3.**
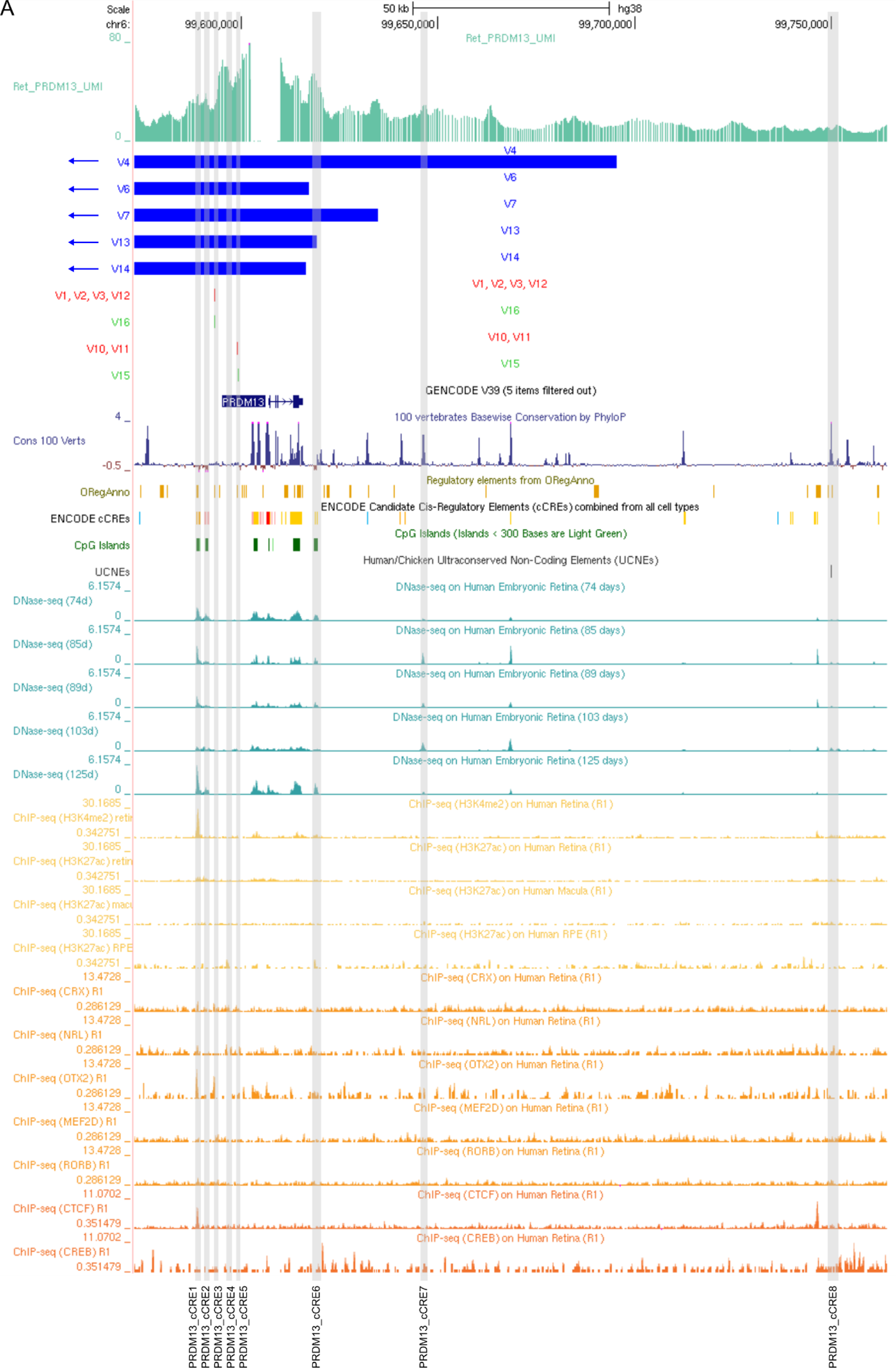
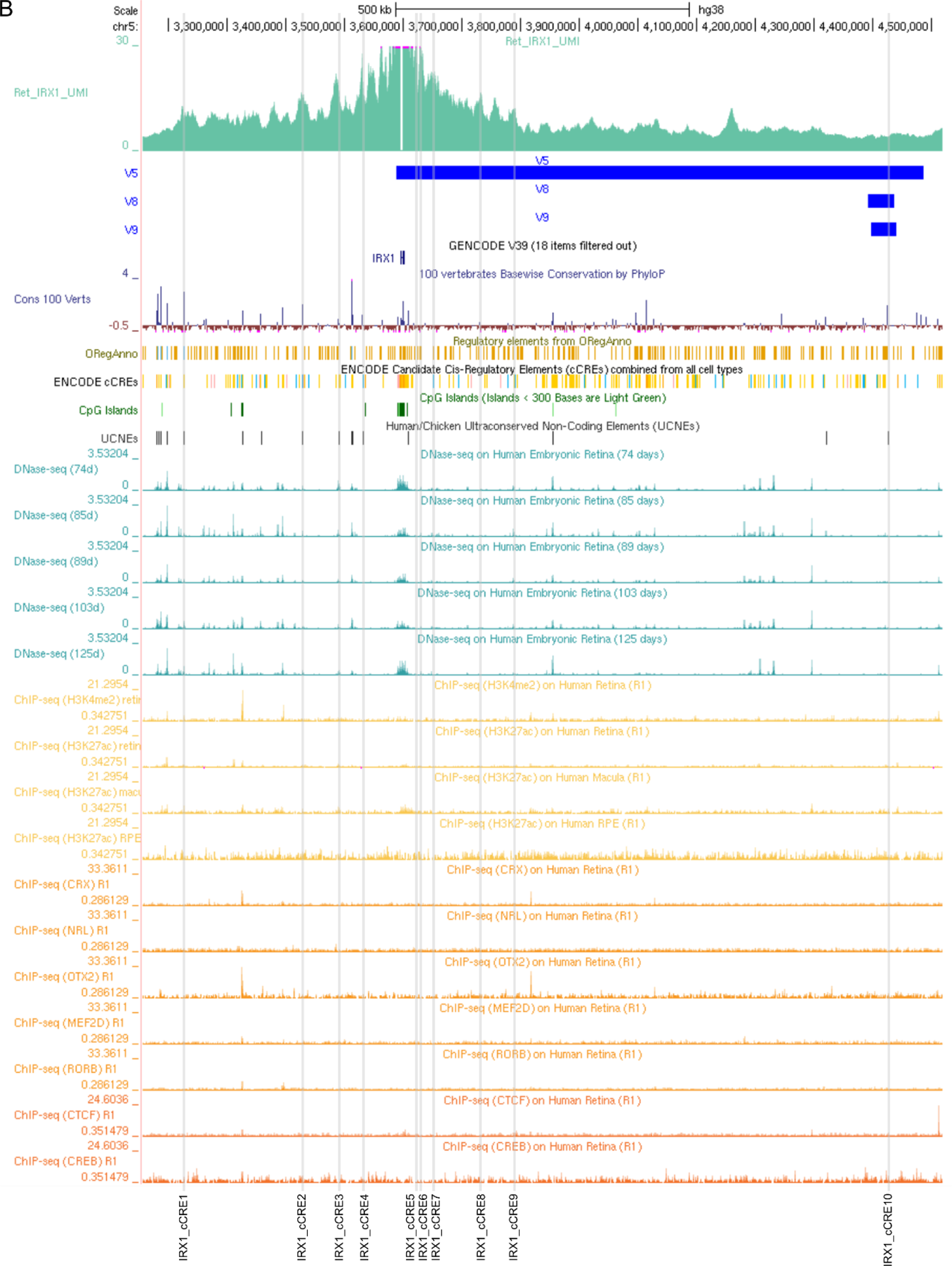
Overview of identified cCREs of the *PRDM13* (A) and *IRX1* (B) loci. The cCREs are obtained after integration of the generated UMI-4C profiles in the retina-specific multi-omics database. The cCREs were defined as non-coding regions interacting with the *PRDM13* or *IRX1* gene promoter while demonstrating overlap with peaks of epigenomics datasets or containing a UCNE or NCMD-associated genetic variants. Known and novel pathogenic SNVs are respectively indicated as red and green bars, while previously reported tandem duplications are indicated as blue bars. cCRE: candidate CRE. UCNE: ultra-conserved non-coding element.

**Figure S4.**
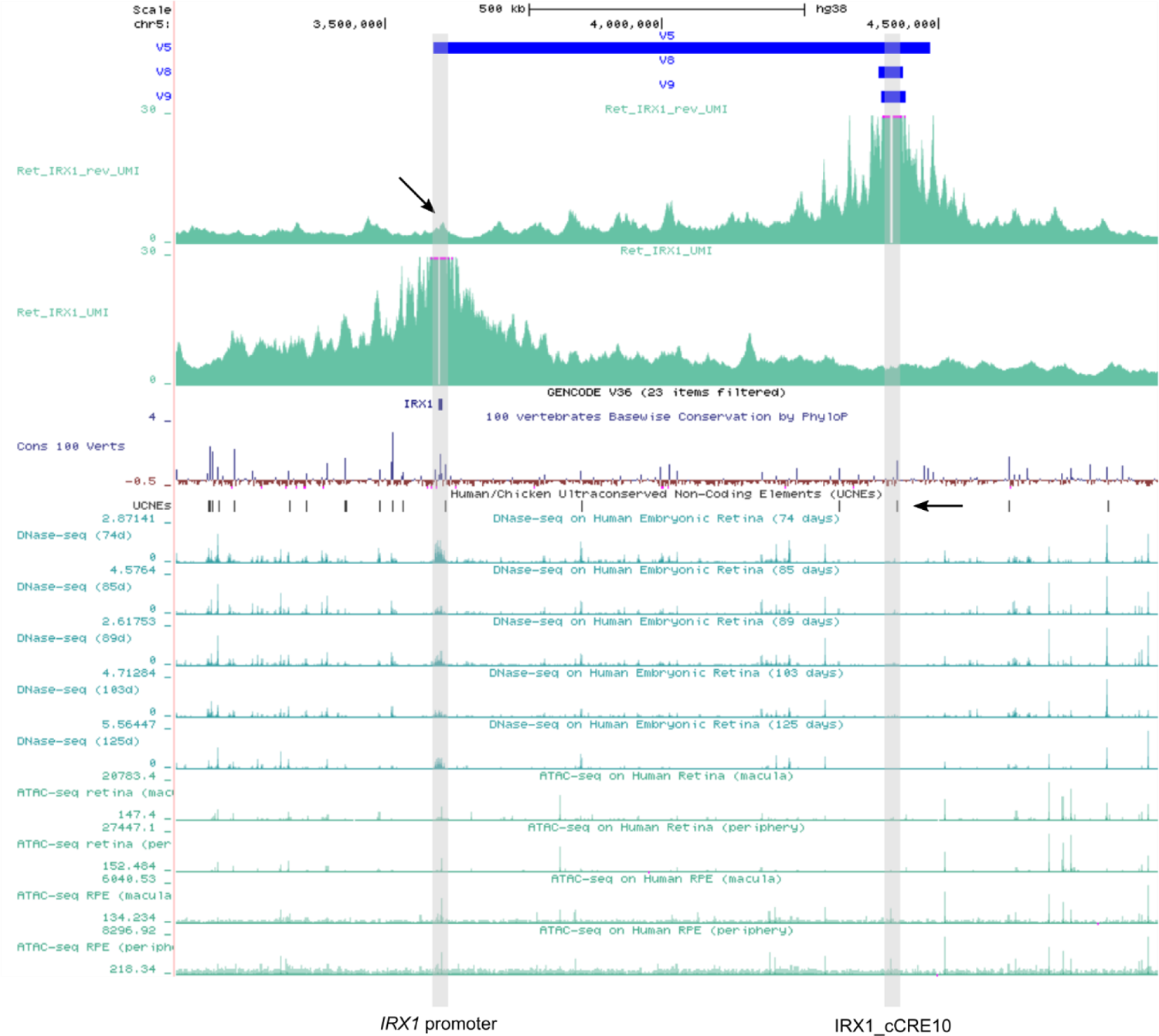
Reverse UMI-4C experiment using IRX1_cCRE10 as viewpoint. The resulting UMI-4C profile suggests an interaction of this region with the *IRX1* promoter, as indicated by the diagonal arrow. However, in the UMI-4C experiment using the *IRX1* promoter as viewpoint, the interaction is less evident. The three tandem duplications are displayed as blue bars. The UCNE located in this region is indicated by the horizontal arrow. UCNE: ultra-conserved non-coding element.

**Figure S5.**
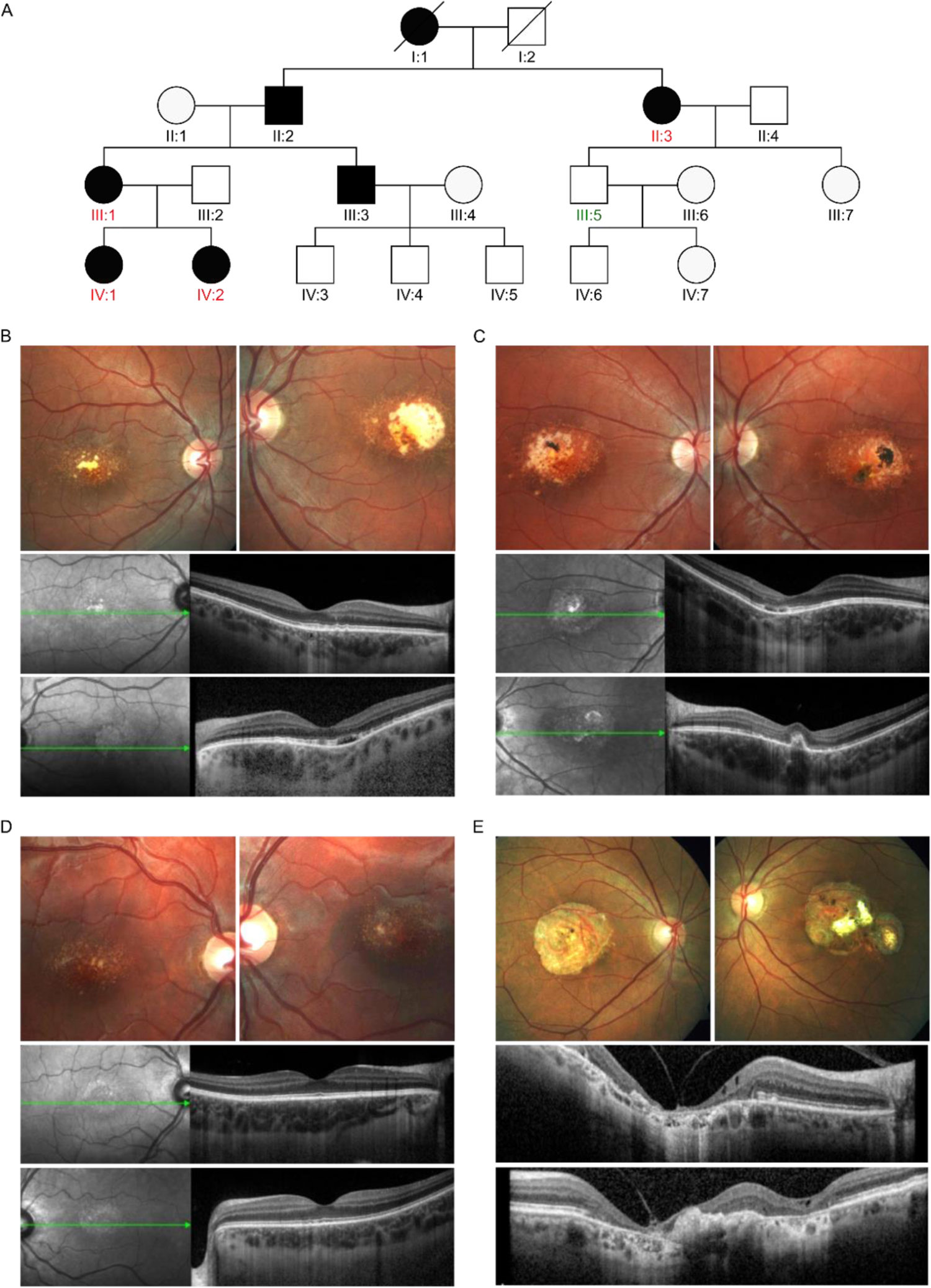
Overview of the genetic and clinical findings in family F1. (**A**) Pedigree of family F1. Individuals that were subjected to genetic testing are colored in red or green, respectively indicating whether the novel heterozygous **V15 variant** (chr6:99599064A>G) is present or not. (**B**) F1-III:1. Fundus BE (top): yellowish-white lesions in the macula, encircled by hypopigmented halos and surrounded by hyperpigmentation. OCT BE (bottom): focal RPE damage and focal neurosensory detachment. (**C**) F1-IV:1. Fundus BE (top): abnormality of macular pigmentation, drusen, focal RPE atrophy, and gross pigment deposits. OCT BE (bottom): focal RPE damage. RE: focal neurosensory detachment. LE: submacular scarring. (**D**) F1-IV:2. Fundus BE (top): mild RPE abnormalities with small macular drusen. OCT BE (bottom): mild RPE abnormalities with small macular drusen. (**E**) F1-II:3. Fundus BE (top): chorioretinal macular atrophy with fibrotic changes more pronounced in the LE. OCT BE (bottom): chorioretinal macular atrophy with fibrotic changes more pronounced in the LE, and vitreofoveal adhesion. This family was previously clinically described by Nekolova *et al.* (2021).^1^ LE: left eye, RE: right eye, BE: both eyes.

**Figure S6.**
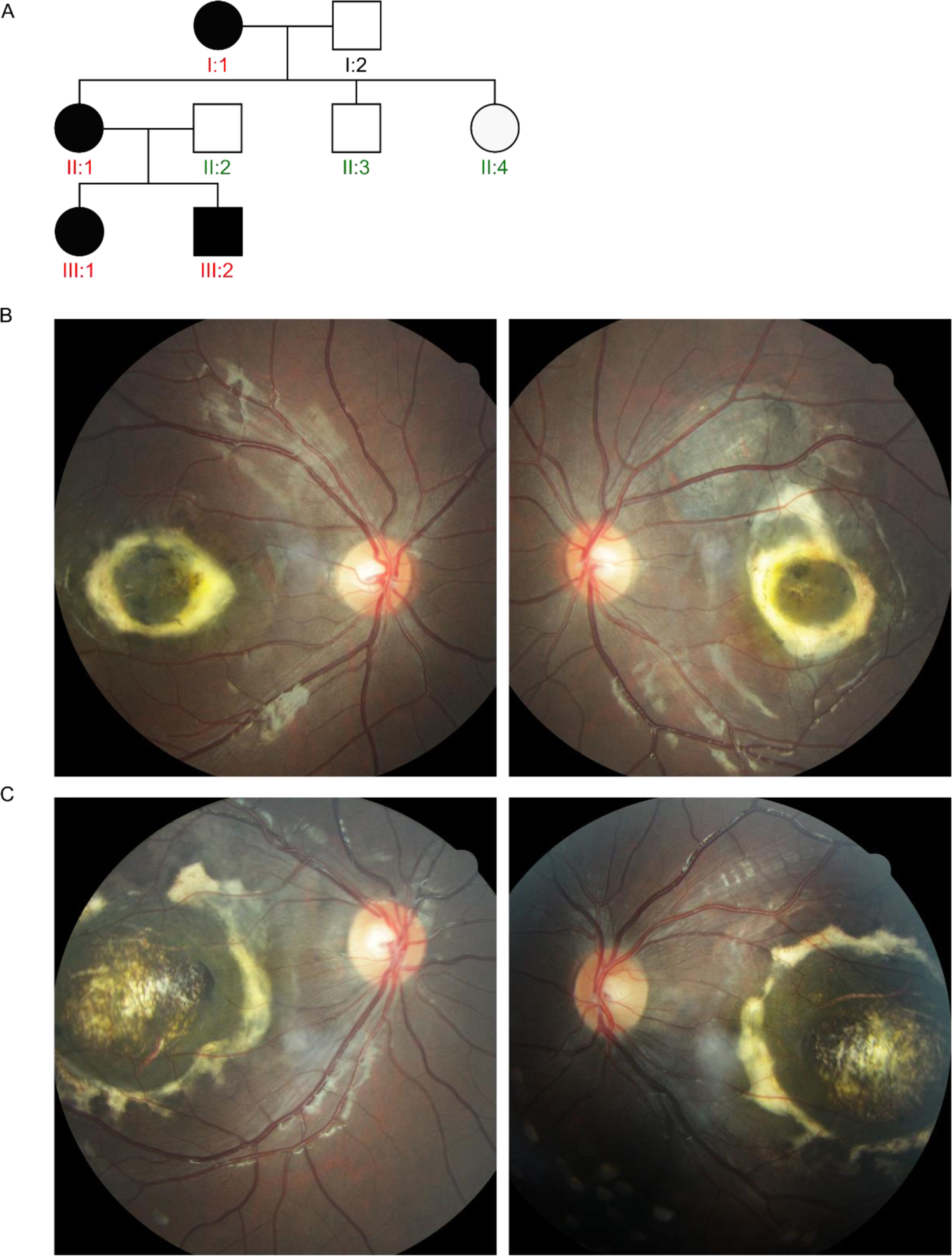
Overview of the genetic and clinical findings in family F2. (**A**) Pedigree of family F2. Individuals that were subjected to genetic testing are colored in red or green, respectively indicating whether the novel heterozygous **V16 variant** (chr6:99593030G>C) is present or not. (**B**) F2-III:1 and (**C**) F2-III:2. Fundus BE: symmetrical coloboma-like macular malformation indicative of grade 3 NCMD. BE: both eyes.

**Figure S7.**
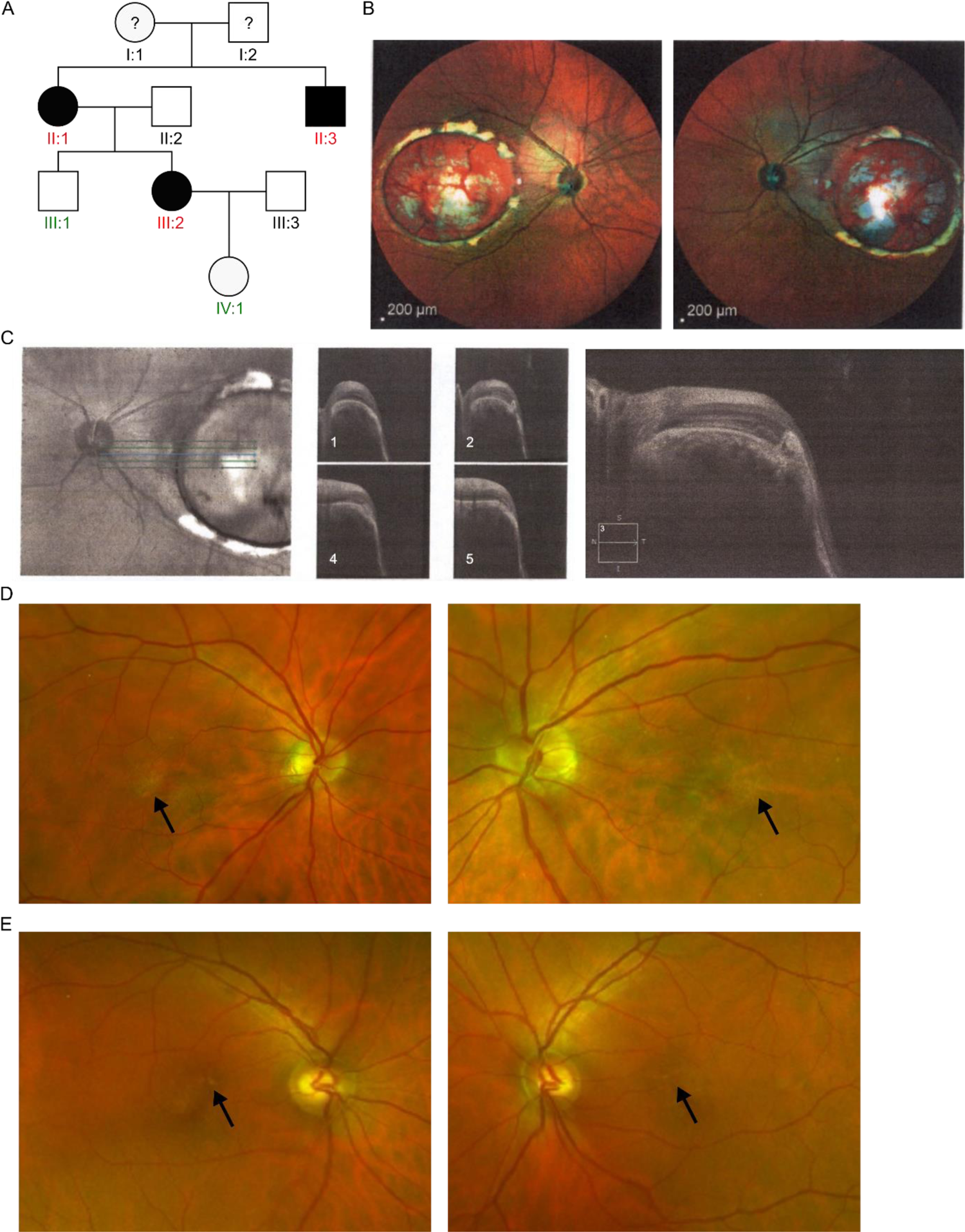
Overview of the genetic and clinical findings in family F3. (**A**) Pedigree of family F3. Individuals that were subjected to genetic testing are colored in red or green, respectively indicating whether the known heterozygous **V1 variant** (chr6:99593030G>T) is present or not. (**B**) F3-III:2. Multicolor (blue, green, and infrared) reflectance images BE: severe bilateral lesions of the RPE affecting the entire macular area, in accordance with grade 3 NCMD (**C**) F3-III:2. OCT LE: excavated lesion with a temporal shelving edge and subretinal fibrosis. (**D**) F3-II:1 and (**E**) F3-II:3. Fundus BE. Mild bilateral yellow specks in the central macula, compatible with grade 1 NCMD are indicated by an arrow. LE: left eye, RE: right eye, BE: both eyes.

**Figure S8.**
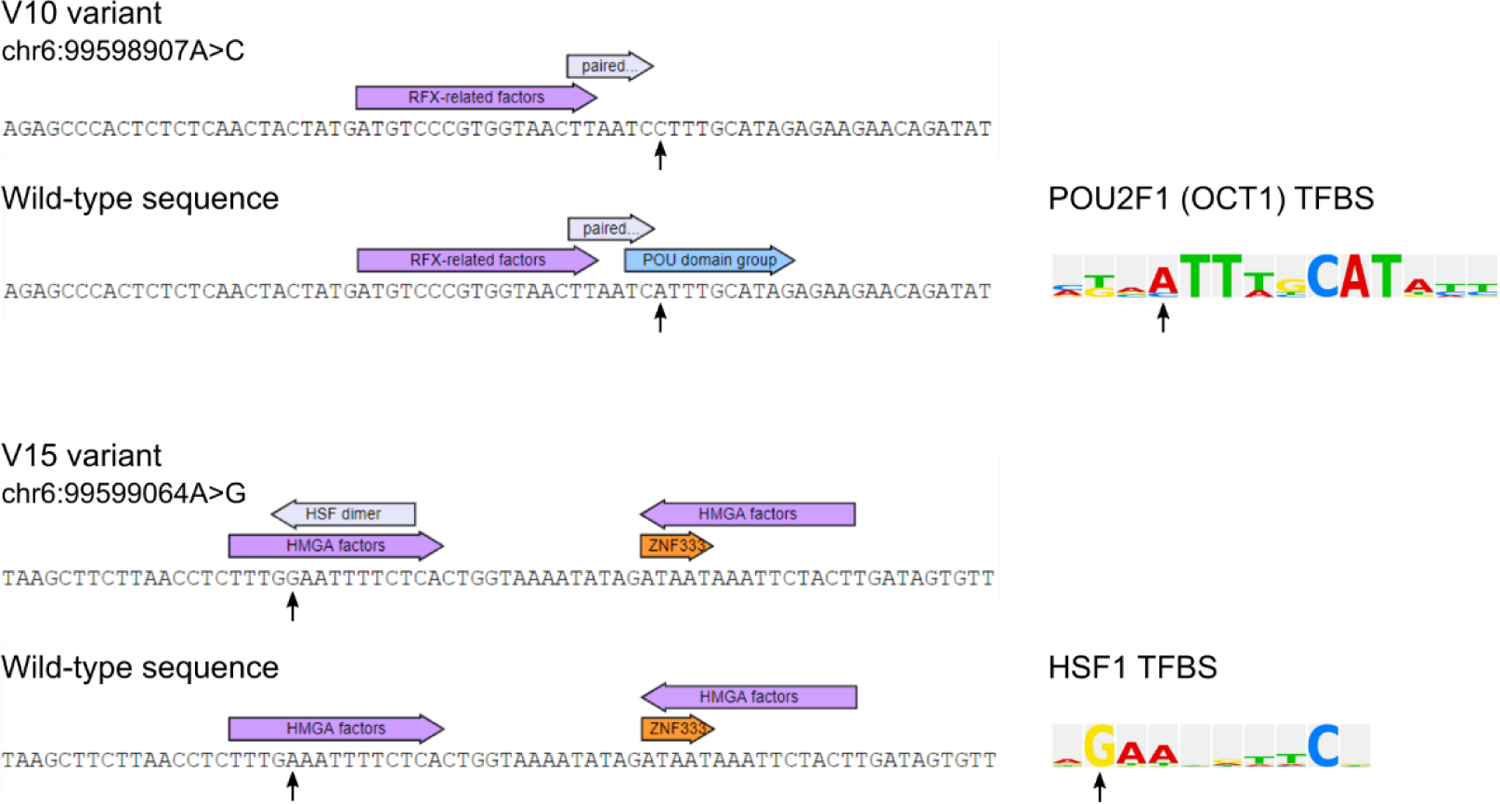
Visual representation of the results from the *in silico* SNV analysis using TRANSFAC. The A to C nucleotide change of the V10 variant is predicted to result in the loss of an POU2F1 TFBS, while the A to G change of the V15 variant results in the predicted gain of an HSF1 TFBS. The nucleotide changes are indicated by an arrow, both in the human DNA sequence and in the binding motif of the transcription factors. TFBS: transcription factor binding site.

**Figure S9.**
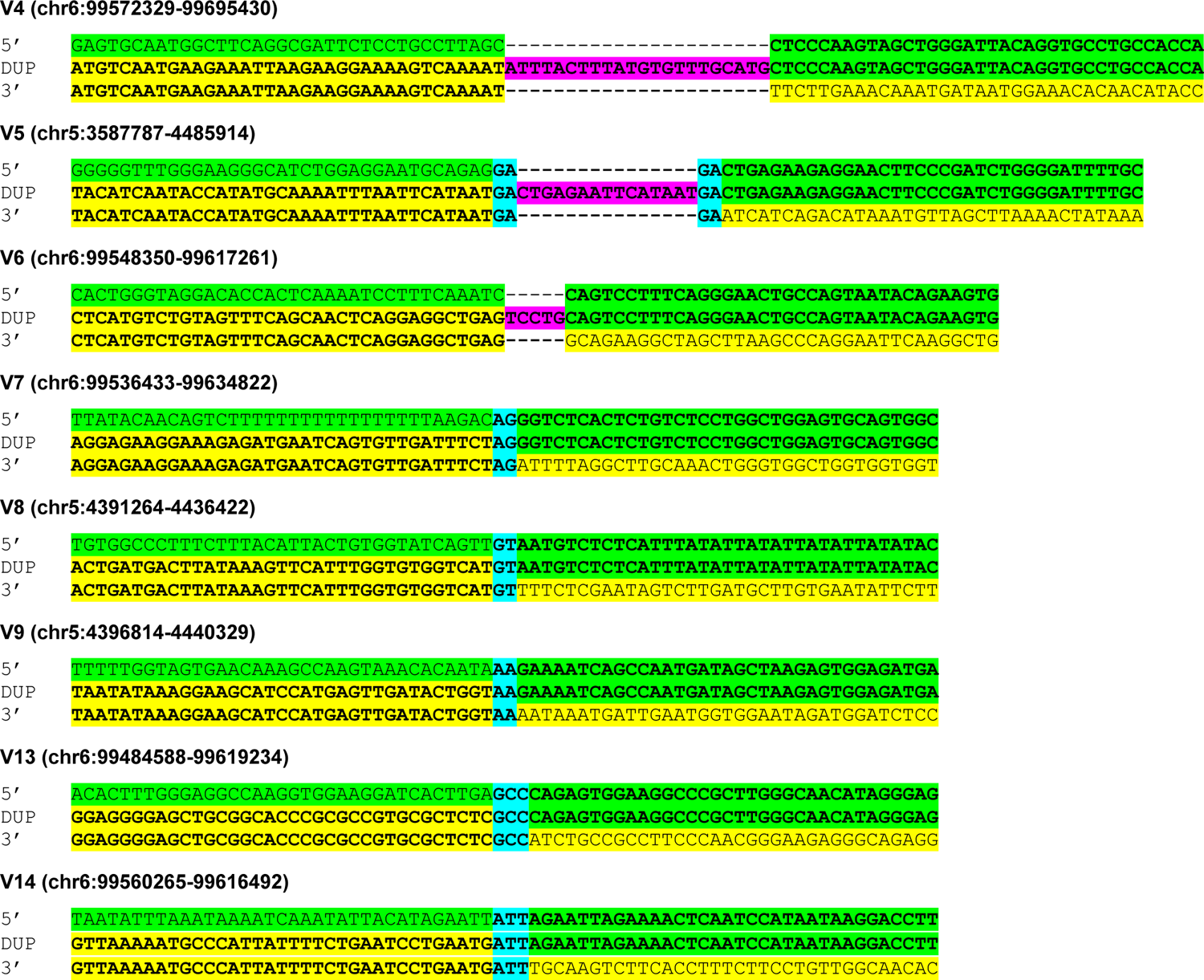
Visualization of the joint point of the eight previously reported NCMD-associated tandem duplications. Inserted bases and microhomology are respectively indicated in pink and blue when present.

**Figure S10.**
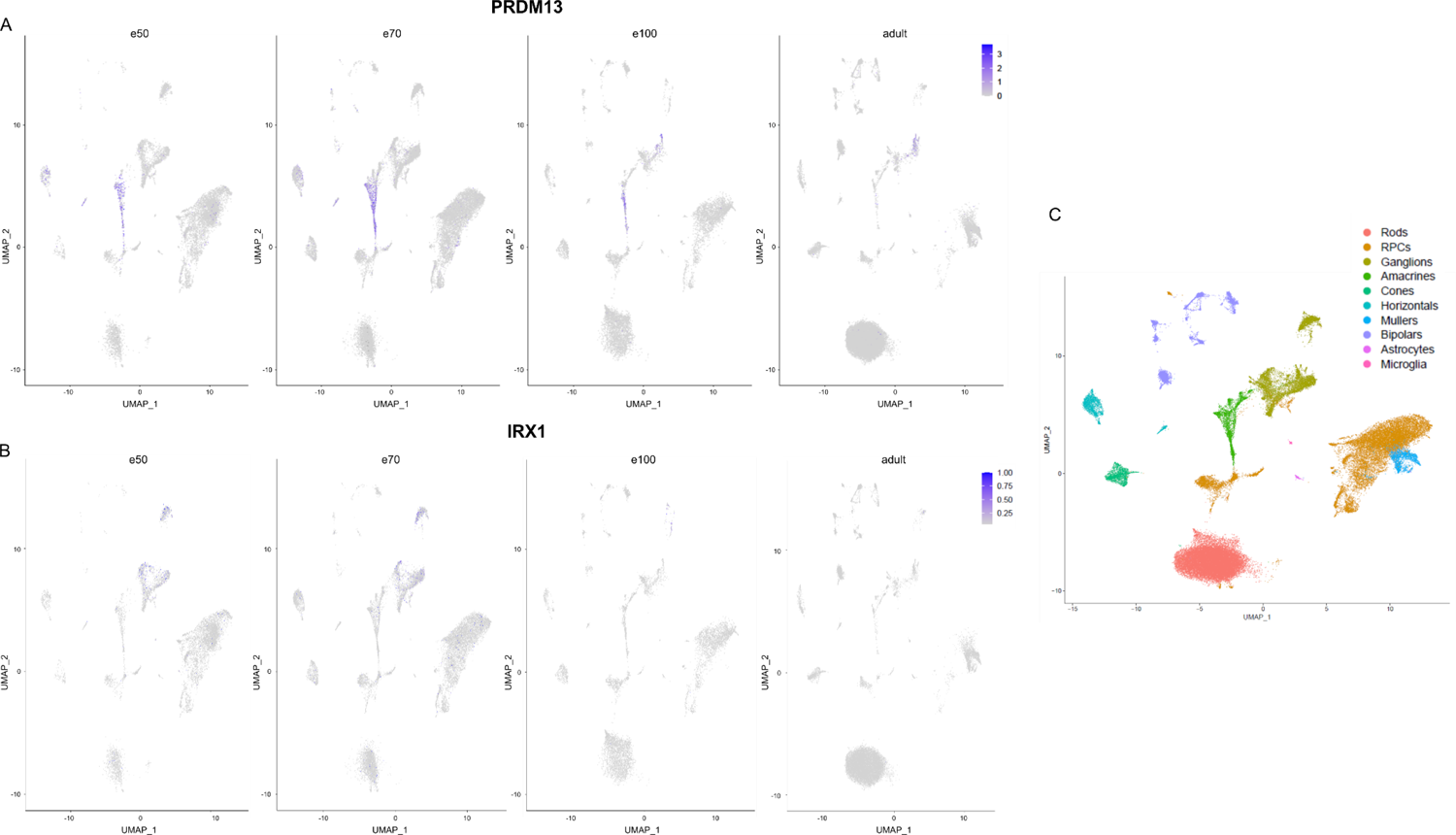
Single-cell transcriptomic analysis of developing human neural retina. UMAP plot showing expression of (**A**) *PRDM13* and (**B**) *IRX1* for four different time points (e50 = d53 and d59, e70 = d74 and d78, e100 = d113 and d132, adult = 3 samples) Plots are scaled to the maximum expression of *PRDM13* and *IRX1*, respectively. (**C**) Annotation of the ten transcriptionally distinct clusters.

**Figure S11.**
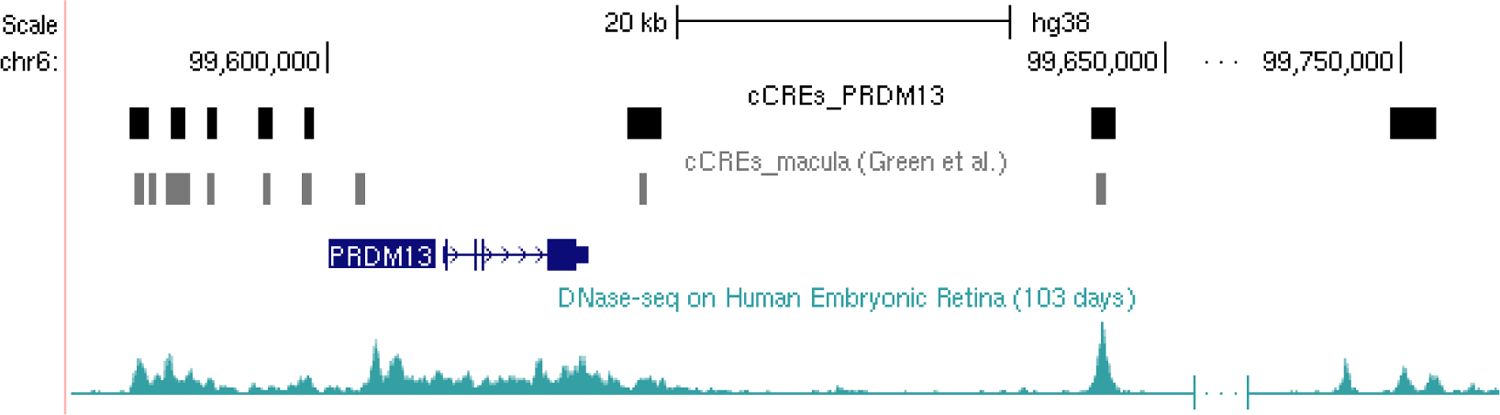
Visualized comparison of cCREs. Comparison of the eight cCREs we identified in the *PRDM13* locus using UMI-4C profiling (black bars) integrated into our retina-specific multi-omics database RegRet with the ten macula-specific CREs identified by Green *et al.* (2021) using the activity-by-contact (ABC) method (grey bars).^2^

